# An innovative approach in identifying a network of priority wetland sites in the East Asian-Australasian Flyway for a new sustainable management investment programme

**DOI:** 10.1101/2025.03.05.641701

**Authors:** Mike Crosby, Shelby Qi Wei Wee, Ding Li Yong, Gary Allport, Sayam Chowdhury, Gan Xiaojing, Ward Hagemeijer, Arne E. Jensen, Duncan Lang, Cynthia Layusa, Yoon Lee, Taej Mundkur, Heejin Oh, Shi Jianbin, Terry Townsend, Doug Watkins, Zeng Qing, Lenke Balint, Stefano Barchiesi, Radhika Bhargava, William Fairburn, Daniel A. Friess, Evelyn Pina-Covarrubias, Karen G. C. Ochavo, Hao Tang, Kelvin S.-H. Peh

## Abstract

The East Asian-Australasian Flyway (EAAF) is widely recognised to be the most threatened of the eight flyways in the world, with wetlands rapidly lost due to land cover change, unsustainable use, and the wider impacts of climate change. The recently established Regional Flyway Initiative aims to bring a network of priority wetlands in the EAAF under improved protection, management, and restoration in 10 Asian countries, while mobilising resources for sustainable agriculture, aquaculture, ecotourism, and other livelihoods for local communities. A major step in the development of this initiative is the identification of priority wetland sites through the application of international criteria, based on modern waterbird count data collated from wetland sites across Asia. Through existing analyses and stakeholder consultations, we short-listed 400 internationally important wetlands as candidate sites for further assessment. Count data of EAAF waterbird species was then assessed against international criteria aligned with the Convention on Wetlands (Ramsar Convention), the EAAF Partnership’s Flyway Site Network and Important Bird Areas and Biodiversity for each site to iteratively identify a subset of priority sites, drawing on newly available species population thresholds. Each site was scored and ranked using a metric (Prioritisation Criterion1) calculated from the proportions of every occurring EAAF species against published population thresholds. We identified a total of 147 wetland sites of high conservation priority across the 10 countries, thereby representing the full suite of wetland landscapes and habitats in Asia, both freshwater and coastal, all critical sites for the conservation of migratory waterbirds in the flyway. At least 34 threatened species, including significant proportions of their global populations are represented in this network of 147 sites. To ensure that conservation opportunities are maximised for species and ecosystem services, there is a need to ensure that selected sites and landscapes are reconciled with the conservation and development priorities of each country, and to evaluate priority sites for their ecosystem services.

## Introduction

The East Asian-Australasian Flyway (EAAF), spanning from the Arctic tundra of central and eastern Russian Federation and Alaska (United States), through East Asia and Southeast Asia to Australasia, represents a vital pathway for over 50 million migratory waterbirds from more than 250 different populations. Notably, this flyway supports at least 34 globally threatened species, making it the most threatened among the eight globally recognised flyways^1,2^. The EAAF is of particular significance to millions of long-distance migratory waterbirds that breed in northern Asia and Alaska and spend the non-breeding season in Southeast Asia and Australasia^3^, and possibly best headlined by the Spoon-billed Sandpiper *Calidris pygmaea*^4^.

Long-distance migratory waterbirds are dependent on connected networks of highly productive wetlands to rest and refuel across their annual cycles. Yet, bird migrations face substantial threats due to the accelerated loss and degradation of wetland habitats^5,6^. This can be attributed to rapid drainage and conversion of wetlands to agricultural, industrial, and urban uses, which have led to the alarming decline of these ecosystems^5,7^. For example, an estimated 1,794.8 km^2^ (29%) of coastal wetlands in the Yellow and Bohai Seas were lost to development between 2000 and 2015 due to conversion for aquaculture and salt pans^8–10^. Historical data suggests that up to 65% of tidal flats in the region have also been lost there over the past five decades^11^. Thus, the remaining wetlands and the migratory waterbirds that they support face a wide range of pressures, including loss of secure roosting sites, illegal hunting, pollution, and the wider impacts of climate change^12–14^. Consequently, an increasing number of migratory waterbird species within the EAAF are in rapid decline^15^, many which have been classified under the IUCN Red List of Threatened Species^16,17^. Some of these species have become flagships for the flyway and the focus for monitoring and conservation actions, including the Spoon-billed Sandpiper^18^, Black-faced Spoonbill *Platalea minor*^19^ and several threatened crane^20^, geese^21^ and duck species^22^.

The implications of wetland loss extend beyond just biodiversity loss, impacting the livelihoods of local communities living in and around wetlands, and the local economies. Fisheries and vital ecosystem services are in danger of collapse and ecological disasters are increasing, with concomitant implications for human communities. For example, wetland loss as a result of conversion to farmland in the Sanjiang Plain in northern People’s Republic of China (PRC) (which includes the Sanjiang National Nature Reserve) has reduced ecosystem service value by approximately $57.46 billion over the past six decades^23^. Therefore, the stakes are very high, including financial loss to the fisheries sector and the potential financial damage and loss of coastal cities, towns, and lands^24–26^.

Urgent long-term and large-scale action is required to address this ecological and socio-economic crisis, thereby securing not only the integrity of wetland sites and landscapes in the EAAF, but also the wellbeing of local human populations and the adaptation to climate change. The Asian Development Bank (ADB), BirdLife International and the EAAFP launched the Regional Flyway Initiative (RFI) in October 2021^27^, a regional initiative aimed at strengthening the management of wetlands of high conservation priority across 10 Asian countries. A major goal of this transboundary initiative is to enhance the conditions of a network of important wetlands that supports migratory connectivity, contributing to the protection of migratory waterbirds. This initiative is also expected to yield measurable co-benefits for local communities, including ecosystem services, economic development, green infrastructure, and climate change mitigation and adaptation^28^.

A major challenge in biodiversity conservation and natural resources management is the efficient allocation of scarce resources to address complex environmental problems^29^. Often a process of ‘conservation triage’ is necessary to ensure that resources are allocated to where they can make the largest impact, for instance the conservation of landscapes with high species richness or threatened species, potentially by accounting for the costs and benefits, and the likelihood of success^29–31^. Conservation prioritisation therefore forms a critical part of conservation decision-making to maximise impact by mobilising resources to sites, species, and ecosystems that are most threatened. However, prioritising sites/landscapes and species for conservation interventions can be challenging if data used in the prioritisation process is limited or is collected in a non-standardised way^32^. Moreover, information used for prioritisation such as conservation costs and anthropogenic threats may be highly uncertain or difficult to obtain^33^. Prioritising sites and landscapes for the conservation of migratory species can also be challenging at large spatial scales because count or abundance-based datasets on migratory species tend to be scarce or are unevenly collected (variable sampling effort) because of the different, and differing methodologies used to survey species.

Numerous approaches to prioritising sites and landscapes for biodiversity conservation exist and can range from simple counts of species richness at a site to complex mathematical and optimisation approaches based on the representation of sets of species and biodiversity/ecological features in a network of sites^31,34^, and how sites can complement others under the framework of systematic conservation planning^30^. An important consideration in conservation prioritisation is the principle of irreplaceability. Irreplaceability can be defined as the potential and relative contribution of a site to a wider conservation goal, and considers the extent to which options for a representative network of sites are lost, if one particular site within the network is lost^35^. Therefore, sites and landscapes can be regarded as more irreplaceable if they are considered to represent a high proportion of the conservation features (i.e. species, biomes) present in an area considered.

This paper aims to outline the quantitative criteria and methodology used to identify the priority sites earmarked for RFI intervention. The process entails compiling an extensive candidate list of wetland sites of perceived high importance across 10 Asian countries, followed by systematic scientific assessments to pinpoint high-priority sites. The identification of these high-priority sites is standardised by rigorous scientific principles (e.g., irreplaceability) and evidence-based criteria, ensuring credibility and effectiveness. This would serve as specific guidance for assessing site conservation opportunities. We then test the selected priority sites for association with wetland types (coastal versus inland wetland), protection status (protected versus unprotected area) and levels of human development; these associations can have implications for the wetland policy development and site management^36^. The implications and lessons gleaned from our analyses can offer broad insights and guidance for the development of other regional initiatives and conceptual frameworks for the conservation of migratory species and wetlands at the regional scale, for instance frameworks in the Americas and Europe/Africa.

## Methods

### Wetland site prioritisation programmes in the EAAF

To prioritise wetlands of high conservation importance, we evaluated and used internationally accepted criteria adopted by three long established government-endorsed programmes to prioritise wetland sites for conservation (Table 1). These are:

1. The *Ramsar Convention on Wetlands of International Importance*^37,38^ provides a framework for international cooperation for the conservation and wise use of wetlands. Contracting parties of the Ramsar Convention designate suitable wetlands that have met its criteria^39^ for inclusion in the list of Wetlands of International Importance, commonly known as Ramsar sites. All 10 RFI countries are contracting parties to the Convention and had designated a total of 148 Ramsar sites (as of 12 September 2023).
2. The *East Asian-Australasian Flyway Partnership Flyway Site Network*^40^ is a semi-formal network of wetland sites, typically nominated by government partners after having been identified for their international importance to migratory waterbirds in the EAAF. The East Asian-Australasian Flyway Partnership (EAAFP) is an informal and voluntary initiative established to protect migratory waterbirds, their habitats, and the livelihoods of people dependent upon wetlands, and consists of 39 Partner organisations including 18 national governments. The governments of nine of the RFI participating countries are EAAFP Partners and they have collectively designated a total of 49 Flyway Network sites (as of 12 September 2023)^40^.
3. The BirdLife International Partnership’s *Important Bird and Biodiversity Areas (IBAs) Programme*^41,42^ is an global initiative that has identified and documented more than 13,000 Important Bird and Biodiversity Areas (IBAs) worldwide, which are places of global significance for the conservation of birds and other biodiversity based on a measurable set of criteria. IBAs are typically also Key Biodiversity Areas (KBAs)^43^. In total, 465 wetland IBAs and KBAs have been identified to date in the 10 RFI participating countries^44,45^.

**Table 1.** Internationally recognised designations of wetland sites in the Regional Flyway Initiative (RFI) countries from the Ramsar Convention on Wetlands of International Importance, the East Asian-Australasian Flyway (EAAF) Partnership Flyway Site Network, and the BirdLife International Partnership’s Important Bird and Biodiversity Areas (IBAs) Programme (as of 31 March 2023).

| **RFI**<br>**Country** | **Government endorsed programme** |  |  |
| --- | --- | --- | --- |
|  | **Ramsar sites** | **EAAFP Network Sites** | **Wetland IBA/KBA** |
| Bangladesh | 2 | 6 | 9 |
| Cambodia | 5 | 1 | 35 |
| PRC | 82 | 20 | 221 |
| Indonesia | 7 | 2 | 71 |
| Lao PDR | 2 | N/A | 15 |
| Malaysia | 7 | 1 | 16 |
| Mongolia | 11 | 11 | 39 |
| Philippines | 8 | 4 | 17 |
| Thailand | 15 | 3 | 17 |
| Viet Nam | 9 | 1 | 25 |
| Total | 148 | 49 | 465 |

A set of nine quantitative criteria has been developed under the Ramsar Convention to identify Wetlands of International Importance, including the four that are relevant to waterbirds (for details see Table 2). Criteria 2, 5 and 6 are used by the EAAFP to identify Flyway Network Sites for migratory waterbirds in the EAAF, and Criteria 2 and 6 have been adopted by BirdLife International to identify IBAs for globally threatened and congregatory waterbirds worldwide.

**Table 2.** The criteria for identifying Wetlands of International Importance (Ramsar Sites) that are relevant to waterbirds (four of the nine Ramsar criteria; see Ramsar, 1971).

|  |
| --- |
| **The Ramsar Sites Criteria** |
| **Criteria based on species and ecological communities** |
| **Criterion 2:** A wetland should be considered internationally important if it supports Vulnerable, Endangered, or Critically Endangered species or threatened ecological communities |
| **Criterion 4:** A wetland should be considered internationally important if it supports plant and/or animal species at a critical stage in their life cycles or provides refuge during adverse conditions |
| **Criteria based on waterbirds** |
| **Criterion 5:** A wetland should be considered internationally important if it regularly supports 20,000 or more waterbirds |
| **Criterion 6:** A wetland should be considered internationally important if it regularly supports 1% of individuals in a population of one species or subspecies of waterbird |

The 1% population thresholds used under Ramsar Criterion 6 are available from Wetlands International through the Waterbird Populations Portal^46^ (WPP), an online, open-access database which provides current and historic population estimates and 1% population thresholds and population boundary maps for definable biogeographic populations of all waterbird species worldwide^47^. We used the new population thresholds published in the East Asian - Australasian Flyway Conservation Status Review^1,47^ for species for prioritising sites, together with their definitions of migratory populations (which exclude some East and Southeast Asian waterbird populations judged to be non-migratory), and which is possibly the most authoritative source of information on waterbird populations in the region of interest.

### Analysis of potential Flyway Network Sites in the EAAF

Jaensch^48^ conducted a flyway-wide assessment of wetland sites and their relative importance to migratory waterbirds at the regional scale to help guide the development of the EAAFP’s Flyway Sites Network. This study compiled a large dataset of waterbird count data from localities throughout the EAAF (a substantial proportion collected through the annual Asian Waterbird Census coordinated regionally by Wetlands International) and applied the following three site prioritisation methodologies. Jaensch’s^48^ analysis identified over 1,000 potential Flyway Network sites that have met the Ramsar/Flyway Network Site criteria, including 467 sites in the 10 RFI participating countries based on the following,

- Prioritisation Criterion 1 (PC1): Derived from the proportion of total size of internationally important populations (those that regularly support 1% of the individuals in a population of a species and meet Ramsar Criterion 6) which had been recorded at the site, summed across all migratory waterbird populations listed for the site in the project dataset
- Prioritisation Criterion 2 (PC2): Number of populations at 1% (or in some cases 0.25%) level (those that meet Ramsar Criterion 6)
- Prioritisation Criterion 3 (PC3): Number of threatened populations: IUCN Red List categories Critically Endangered, Endangered or Vulnerable (those that meet Ramsar Criterion 2)

Jaensch^48^ concluded that Prioritisation Criterion 1 (PC1) was the most useful metric as a relative measure of the contribution of each site to the conservation of migratory waterbirds in the flyway. Xia *et al.*^49^ prioritised wetland sites of importance to migratory waterbirds in the PRC’s coastal wetlands, and used a similar prioritisation methodology to Jaensch’s^48^ PC1, represented with the formula:

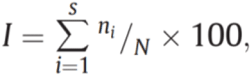

where *ni* denotes the population of *i*th species of waterbirds at the survey site, *N* denotes the population of *i*th species globally or throughout the flyway, and s denotes the number of species at the survey points (see Figure 1).

**Figure 1.**
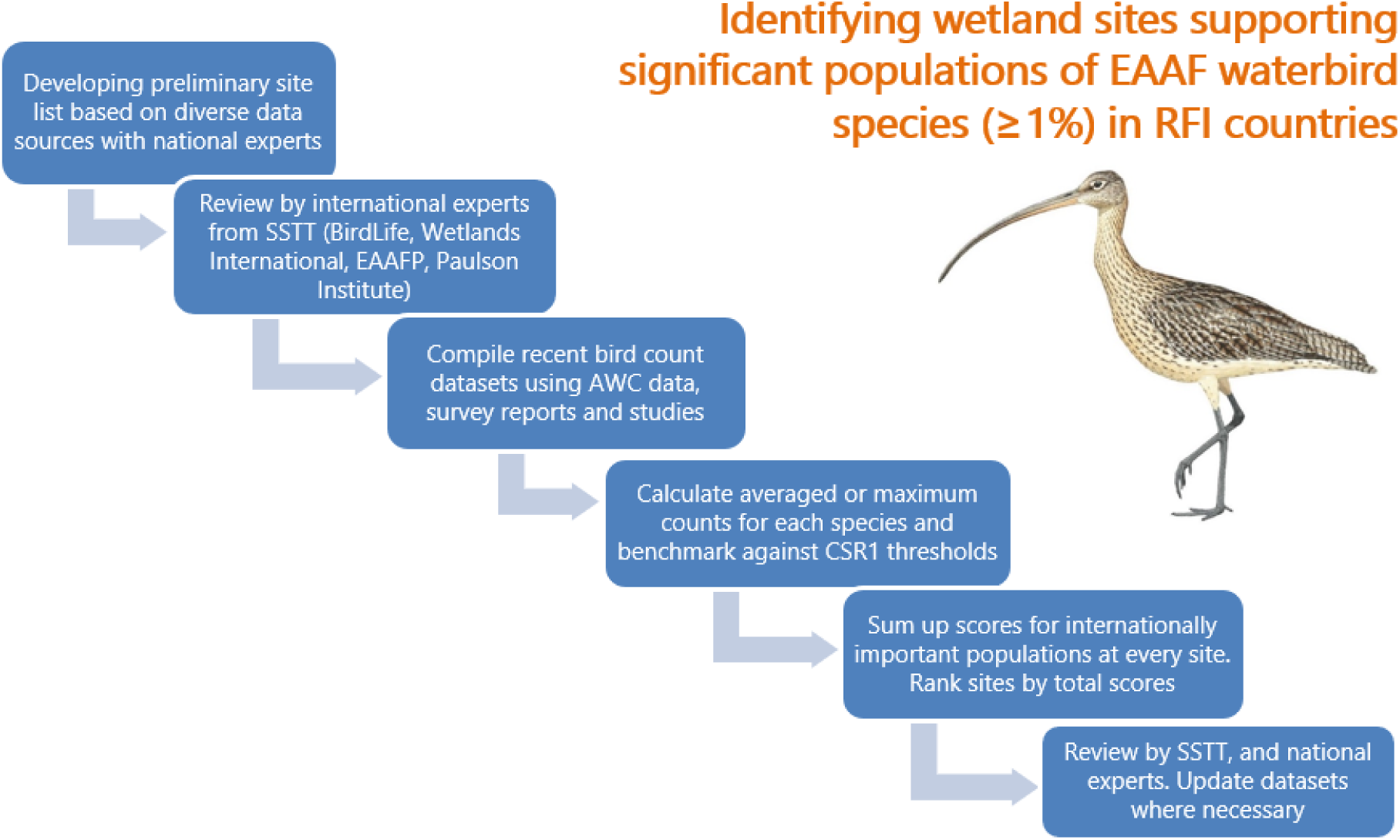
Regional Flyway Initiative (RFI) site selection framework for the East Asian-Australasian Flyway (EAAF): key steps in the site prioritisation process. Members of Site Selection Task Team (SSTT) were coauthors of this paper. AWC stands for Asian Waterbird Census. CSR1 was the East Asian-Australasian Flyway Conservation Status Review.

### Sources of data on migratory waterbirds and wetland sites

We compiled count data on wetland sites from 10 Asian countries on the numbers of migratory waterbirds reported at these sites (for details see Supplementary Tables 1 and 2). The main sources of data are outlined below.

#### Site prioritisation analyses

The c.400 bird count localities listed by Jaensch^48^ for the 10 RFI participating countries formed the foundation for the RFI sites prioritisation analysis and provided a large (but relatively old) database of waterbird count records. Additional data on the definition and locations of sites and waterbird abundances (maximum counts) was compiled from flyway-wide analyses of EAAF wetland sites by MacKinnon *et al.*^24^ and important shorebird sites by Conklin *et al.*^50^, and the analyses of priority wetlands in the PRC by Paulson Institute^51^, Xia *et al.*^49^, Zhang *et al.*^52^ and Duan *et al.*^10,53^.

#### Internationally and nationally designated sites

Data on Ramsar sites, UNESCO World Heritage Sites^54^, EAAFP Flyway Network sites and IBAs was compiled, including boundary maps, site information sheets and reports with waterbird count data. The boundary maps of protected areas, where available, were downloaded from the World Database on Protected Areas^55^ (WDPA).

#### Asian Waterbird Census (AWC)

The AWC^56^ gathers information annually on waterbird populations at wetlands in the region during the non-breeding period of most species (January), as a basis for the evaluation of sites and monitoring of populations^57^. Together with the historical data published from the AWC from 1987 to 2015, this formed the largest source of data for our analyses^57,58^.

#### National and flyway-wide waterbird monitoring schemes

Data on internationally important counts of migratory waterbirds at the site level was obtained from diverse sources. In the PRC, the China Coastal Waterbird Census^59,60^ provided the main source of count data for species at the site level. Additionally, site-level count data was also compiled based for non-breeding waterbirds in the Central and Lower Yangtze river basin^61,62^, Northeast China^63,64^ and in the Yellow Sea-Bohai based on large scale counts^65^.

Data on sites across the EAAF was also compiled based from field projects under the Global Flyway Network^66^ (e.g. Hassell *et al.*^67^, Piersma *et al.*^68^, the International Black-faced Spoonbill Census^69^, the Spoon-billed Sandpiper Winter Census^70^, and other waterbird survey projects funded by the EAAFP and conservation organisations such as BirdLife International, Wildlife Conservation Society (Cambodia and Lao People’s Democratic Republic [PDR]), Manfred Stiftung/Eksai foundations (Indonesia) and IUCN (Lao PDR)).

#### Peer-reviewed scientific papers and survey project reports

A systematic literature search was conducted to identify peer-reviewed articles that held count data on migratory waterbirds and wetland sites in the EAAF. This covered every issue of six relevant bulletins and journals from 2010 to 2021; they include *Wader Study*, *Stilt*, *Bird Conservation International*, *Forktail*, *BirdingAsia* and *Kukila*.

#### Expert opinion and other sources of data

We consulted the national experts (see Acknowledgements) who provided information on the relative importance of many of the candidate priority sites and advised on other sources of grey literature on wetland sites and waterbirds. We also consulted other sources of data, notably waterbird counts at potentially important sites downloaded from the eBird website^71^, which are validated by national experts, several of whom are authors of this paper (TM, AJ).

### Definition of wetland sites and their boundaries

While there have been many initiatives to assess the abundance of waterbirds in the region, there is no standardised (consistent) list of sites and their names at the international/regional level, nor are there broadly agreed demarcations of wetland site boundaries^48^. We worked with national experts to address this issue by reviewing the definitions of candidate priority sites in the 10 participating countries and exploring how waterbird counting locations could be (hierarchically) clustered together, with the aim of defining areas that are potentially suitable site management units (following guidelines developed by BirdLife’s IBA Programme, and the AWC Guidelines^72^ on count site mapping developed by Wetlands International. An associated issue is that many wetlands in the East Asian-Australasian Flyway have been affected by land claim, wetland conversion to urban, industrial, and agricultural land (e.g., Caofeidian wetlands in the PRC, Saemangeum in the Republic of Korea and Manila Bay, Philippines), pollution and other forms of degradation^24^. We have considered how this habitat loss and other threats might have impacted the number of waterbirds, and therefore, affected the PC1 scores for these sites.

### Management and analysis of waterbird count data

We designed a spreadsheet to store and analyse the waterbird count data (Supplementary Table 1). The fields in the spreadsheet include the species and site details, count data, data sources, and species scores. As waterbird count data was extracted from the data sources outlined above, it was entered into the spreadsheet, one record per row (or where suitable data was available, the average of counts of a species over several years). Once the data collection was completed for each country, the spreadsheet was used to calculate the maximum scores assigned to the species recorded at each site in relation to the 1% population thresholds. Scores for all migratory waterbird populations that exceeded the relevant 1% CSR1 thresholds were used to calculate the overall PC1 score for each site.

We considered a range of issues related to site selection and data analysis. The original focus of the RFI was on coastal wetland sites, but two of the participating countries are landlocked (Mongolia and Lao PDR) and the national experts recommended including inland wetlands in the Yangtze and Yellow River basins and Northeast PRC because of the waterbird species that they support, and the potential economic and social co-benefits are different from sites in the relatively prosperous coastal provinces.

At sites which were regularly monitored, we averaged the most recent maximum counts of migratory waterbird species over a three- or five-year period. However, we noted that many sites in the EAAF are irregularly monitored, and in many instances a single maximum count may be used given the paucity of data. In the PRC, we used data from primary sources where available, in particular the China Coastal Waterbird Census (e.g. Bai et al.^59^). Some poorly known wetlands are little surveyed due to accessibility issues and available data on bird counts is dated. In such cases, we relied on expert knowledge of changes in the condition of wetland habitats, and the status and occurrence of migratory species. In general, datasets collected before 2005 were used only in exceptional cases, such as for Dongsha shoals in Jiangsu Province, PRC^73^.

### Developing a network of RFI priority sites throughout the East Asian-Australasian Flyway

We identified a set of priority sites in each of the 10 countries, with the aim of creating a flyway-wide ‘network’ of priority sites. We adopted the same criteria and methodology in all countries except for the PRC and Mongolia where we adopted a threshold PC1 score >10. We had decided to adopt such as an approach as wetlands in the PRC and Mongolia are generally extensive and support high concentrations numbers of (staging) migratory waterbirds, as well as a higher diversity of migratory waterbird species than the countries in the southern part of the EAAF.

### Characterising the RFI priority sites: wetland types, protection status and levels of human development

We classified all prioritised sites according to their wetland types (either coastal or inland), including artificial wetlands such as salt pans and fishponds, which are present in many of the priority sites, and typically provide important feeding and roosting habitats for migratory waterbirds. Coastal wetlands are defined as those found along coastlines and estuaries, characterised by varying salinities due to mixing of seawater and fresh water, creating challenging conditions for most plants; they encompass tidal flats, a vital feeding habitat for a suite of specialised waterbirds, salt marshes and mangroves^74^. Inland wetlands are diverse, occurring on floodplains, depressions, and low-lying areas, with varying water presence; they encompass lakes, rivers, marshes, swamps, and wooded swamps dominated by different types of vegetation^74^.

Also, we classified each priority site as either a protected/partially protected area or unprotected area based on available information on the site. We considered a protected area as a distinctly defined geographic area that is identified, designated, and administered, either through legal methods or other effective approaches, to ensure the long-term preservation of nature, along with its linked ecosystem benefits and cultural values (following IUCN definition^75^).

Lastly, we grouped each priority site according to their country’s human development index (HDI), which is derived from a diverse range of socioeconomic indicators, such as life expectancy, literacy rate, access to electricity in rural areas, GDP per capita, trade activity, homicide rate, multidimensional poverty index, income inequality, internet accessibility, and numerous others^76^. These indicators are synthesized into metric ranging from 0 to 1.0, with 1.0 representing the highest level of human development. The HDI is then categorized into four tiers: very high human development (0.8-1.0), high human development (0.7-0.79), medium human development (0.55-0.70), and low human development (below 0.55).

Countries with low human development tend to have unstable governments, widespread poverty, limited access to health care, poor education, low income, and low life expectancies. We used Chi-square goodness of fit test to analyse if the selected priority sites are associated with a particular wetland type, protection status, or HDI tier. All analyses were performed using R^77^.

## Results

### Outcomes of RFI priority site identification

We designed and implemented a rigorous protocol for wetland selection, as fully detailed in the Method section. A total of 147 RFI Priority Sites were prioritised in the 10 countries (Fig. 2; Table 3; for details see Supplementary Table 3) from the larger pool of wetland sites identified for our analyses. The number of sites per country ranges from three in Lao PDR to 60 in the PRC, and all 10 countries have a range of opportunities to develop site-based investments and hence, contribute to the conservation of migratory waterbirds and management/restoration of wetland habitats, and the maintenance of migratory connectivity within the EAAF. They include 91 wetland sites located on or near the coast (sites dominated by mangroves and other intertidal wetlands), and 56 inland (mostly freshwater) wetlands along major river basins (Table 4).

**Figure 2.**
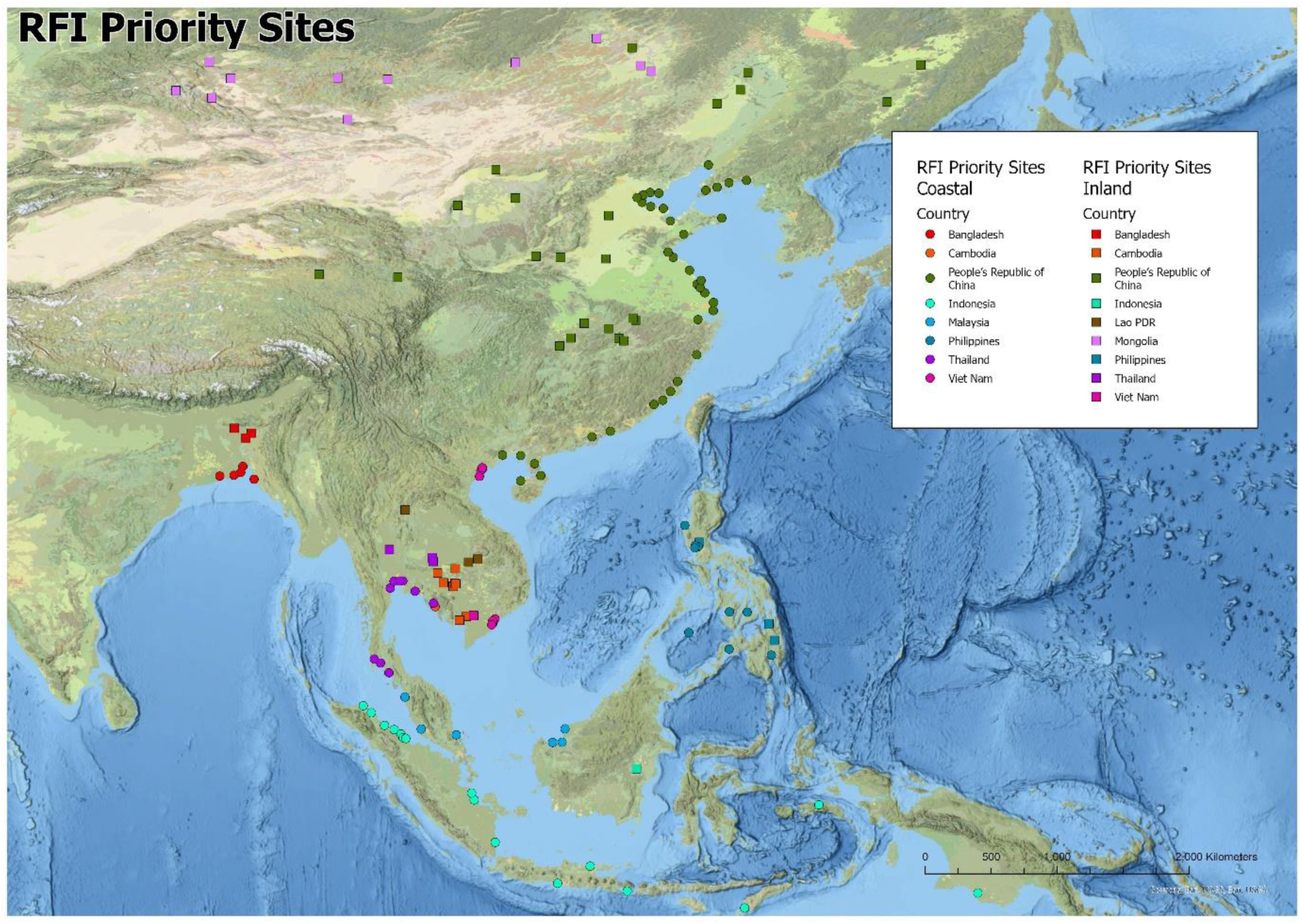
Regional Flyway Initiative (RFI) priority sites.

**Table 3.**
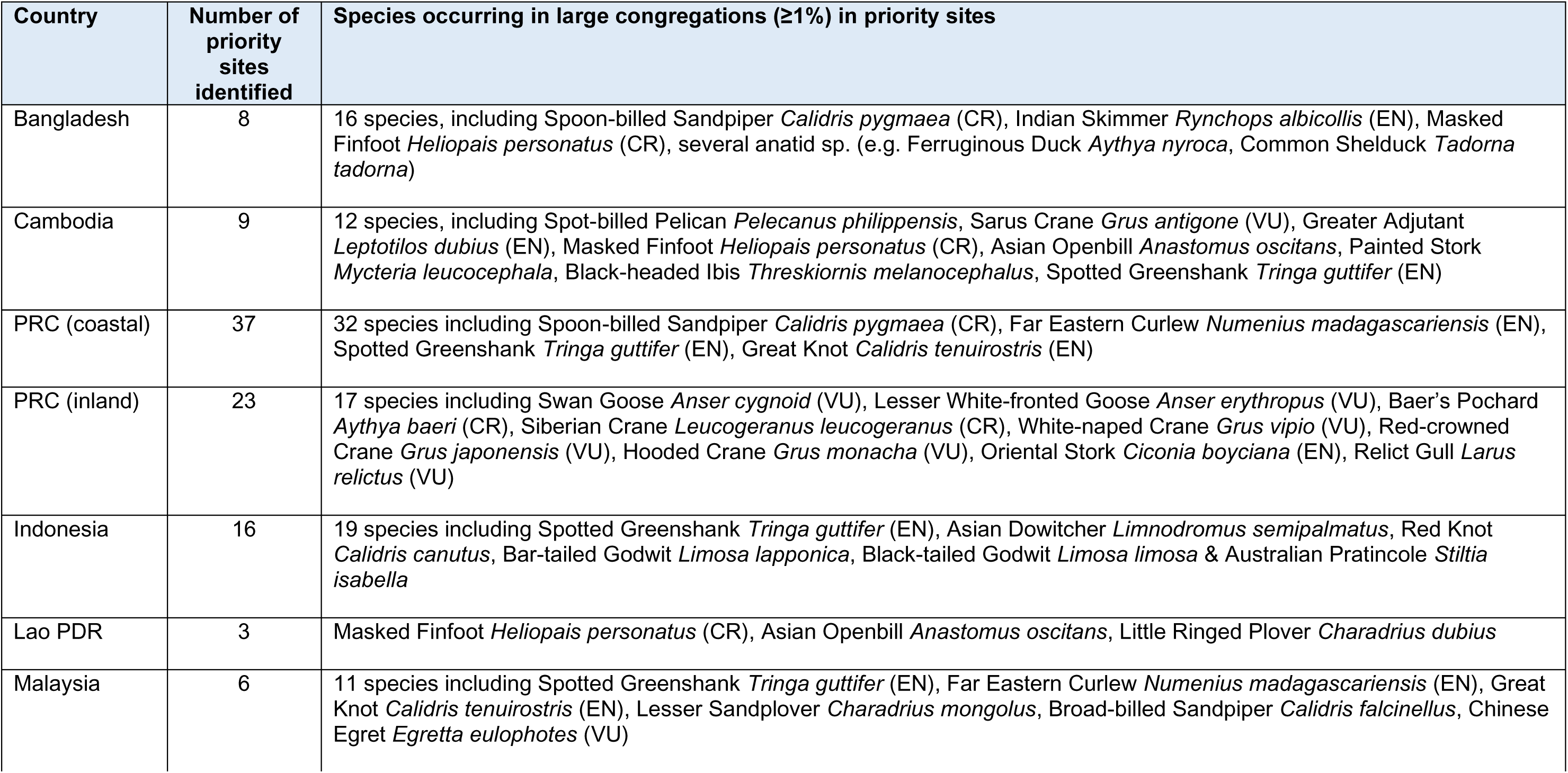
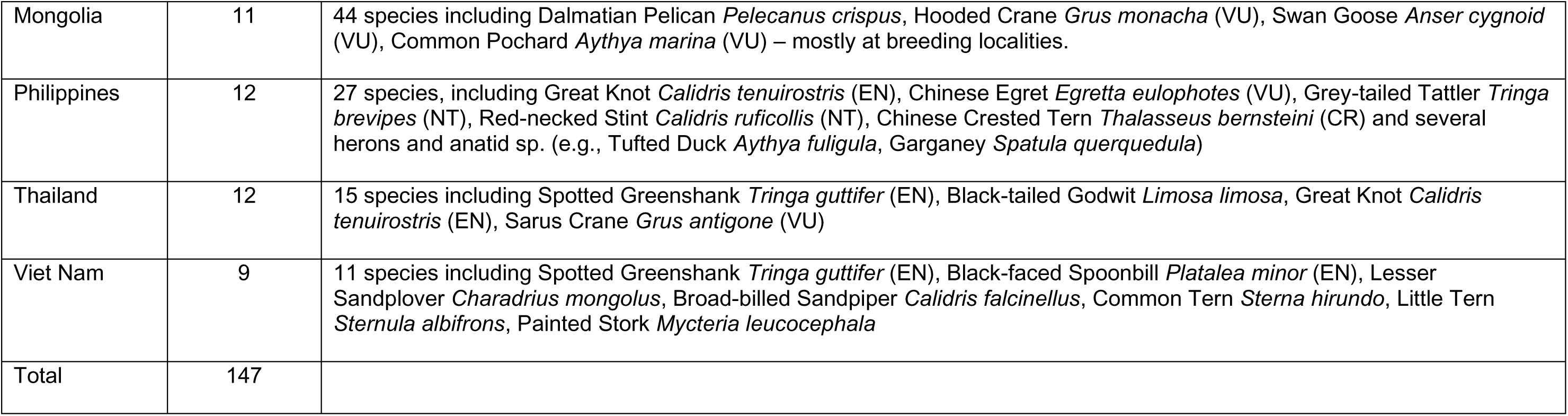
Summary of information on the identified RFI priority wetland sites for the 10 participating countries, including the total number of priority sites identified in each country, number of species that occur in large congregations (≥1%), and IUCN Red List status for globally threatened species (CR = Critically Endangered; EN = Endangered; VU = Vulnerable).

**Table 4.** Priority sites in the 10 Regional Flyway Initiative (RFI) countries.

| Country | Number of RFI priority sites | Wetland type |  | Protection status |  | Human Development Index tier (2021/22) |
| --- | --- | --- | --- | --- | --- | --- |
|  |  | Coastal | Inland | Protected or partially protected | Unprotected |  |
| Bangladesh | 8 | 5 | 3 | 8 | 0 | Medium |
| Cambodia | 9 | 1 | 8 | 7 | 2 | Medium |
| PRC | 60 | 37 | 23 | 52 | 8 | High |
| Indonesia | 17 | 16 | 1 | 8 | 9 | High |
| Lao PDR | 3 | 0 | 3 | 2 | 1 | Medium |
| Malaysia | 6 | 6 | 0 | 2 | 4 | Very high |
| Mongolia | 11 | 0 | 11 | 10 | 1 | High |
| Philippines | 12 | 9 | 3 | 7 | 5 | Medium |
| Thailand | 12 | 9 | 3 | 8 | 4 | Very high |
| Viet Nam | 9 | 8 | 1 | 4 | 5 | High |
| TOTAL | 147 | 91 | 56 | 108 | 39 |  |

Table 3 summarises the data gathered on these sites in the 10 RFI countries and why they were selected. To interpret the analyses in each country, it is important to be aware of the variations in (i) the geographic distributions and abundance of migratory waterbird populations within the EAAF; (ii) the characteristics of the wetlands in different parts of the flyway; and (iii) the availability of data on the numbers of migratory waterbirds that occur at wetland sites, as well as differences in the geographic sizes and topography of the countries. The national waterbird and wetland experts provided guidance on how to factor these into the selection of the priority sites. Caution should be exercised in making comparisons between countries because of variations in distribution of various waterbird species, the characteristics of the wetland sites, and the availability of data.

Additional information on the priority sites is given in Supplementary Table 2, including the site name and province; central coordinates; species that exceed 1% of the EAAF population at the site; IUCN Red List status for globally threatened species (CR = Critically Endangered; EN = Endangered; VU = Vulnerable); outstanding waterbird population (* = exceeding 10% of CSR1 estimate; ** = exceeding 50% of CSR1 estimate); and RFI priority site score.

### Characteristics of the RFI priority sites: wetland types, protection status and human development index

Of the 147 selected priority sites, the distribution between the two wetland types (coastal wetlands = 62%, inland wetlands = 38%; Table 4) was not equal (χ^2^ = 8.33, df = 1, P <0.004). The distribution of the sites between the two protection statuses (protected/partially protected sites = 73%, unprotected sites = 28%; Table 4) was also not equal (χ^2^ = 32.39, df = 1, P <0.0001). The number of sites distributed across countries of three HDI tiers (very high human development = 12%, high human development = 66%, and medium human development = 22%) was also not equal (χ^2^ = 72.53, df = 2, P <0.0001). Most of the sites were associated with countries of either very high or high human development (combined = 78%). None of the countries belonged to the tier of low human development.

## Discussion

### Selection of the RFI priority sites

The EAAF Regional Flyway Initiative aims to prioritise conservation and management interventions for the most important wetland sites for migratory species through a data-driven site selection process to assess internationally important wetlands in 10 Asian countries. In our analyses, we adopted site selection criteria in alignment with the quantitative criteria developed for three long-established, programmes and initiatives for wetland conservation in Asia: the Ramsar Convention on Wetlands, the Flyway Site Network of the EAAFP, and the BirdLife International’s civil-society led IBA Programme. Our method uses an approach similar to an analysis for development of a network of priority wetland sites that Jaensch^48^ conducted for the EAAFP and through the consensus of wetland conservation experts. The selection of priority sites was based on the best available waterbird count data and in consultation with national experts. The number of selected priority sites was more than expected, primarily because more inland sites in the PRC were included on the list than originally planned, following guidance from experts on the importance of these sites and their high potential for conservation investments.

### Site-based approach to conservation

In this study, we focus on the site-based approach as a pivotal tool for biodiversity conservation, recognising its limitations and role within a broader context of conservation strategies^78^. The dispersed nature of migratory species renders the conventional site-based approach insufficient for their effective conservation^79^. For example, many waterbird species covered in this study may be highly dispersed during the breeding season but congregate in large flocks during the migration passage and non-breeding period, which is when they are in habitats that are most threatened in their entire distribution, or are vulnerable to hunting. Conservation of a network of important sites can be an effective approach for migratory species if sites prioritised in the network are of high importance for the species, which can be judged based on high proportions of the whole population being present at the site.

Traditionally, actions to conserve migratory taxa have been associated with a site-based approach, leveraging the concentration of these species at specific wetlands (e.g., Mehlman et al.^80^). However, our site-based approach should be considered one among multiple strategies in the conservation toolbox. Our findings provide for a framework of a threefold approach to migratory species conservation:

1. Highlighting species-specific strategies, particularly for globally threatened birds within flyways (e.g., Black-faced Spoonbill, Spoon-billed Sandpiper) can direct resources toward their conservation. While an individual species focus might not always yield cost-effectiveness (see Lloyd et al.^81^), it could guide conservation investments for sites of importance to many species due to the perceived importance of these species (see Fitzpatrick et al.^82^).
2. Selecting a strategic set of sites that are designed to generate tangible impacts on bird populations^83^ at the range-wide and continental level. The site-based approach remains the cornerstone of our project. However, the historical piecemeal implementation of this approach across flyway has limited its impact. Our targeted site-based network strategy marks a significant departure from previous efforts and holds promise for population-level changes^84^.
3. Emphasising the need for policy-driven strategies. These encompass interventions at broader scales, with agricultural policies and investments in landscape-altering platforms akin to the Farm Bill in the United States or the Structural Funds and The Birds Directive^85^ in the European Union. Such policies can address the conservation needs of migratory species that transcend site-specific boundaries (e.g., Great Lakes Commission^86^).

The synergy among these three strategies is essential for effective biodiversity conservation. Our study challenges the notion that a single approach can suffice, emphasising the importance of an integrated conservation strategy. Despite the limitations of the site-based approach, it is vital to recognise its role within a larger ecological network. Species losses are driven by various factors and, although the site-based approach can mitigate localised threats, it may not address global challenges such as climate change. However, our study pioneers a comprehensive, large-scale implementation of the site-based approach, aiming to contribute to reversing the declining trends observed in critical regions, such as the Yellow Sea.

### Limitations in knowledge and data on waterbird populations

Using data from diverse sources for our analyses, we acknowledge that there are considerable variations and uncertainties in the count data of shorebirds from surveyed sites across 10 countries. Some wetlands in the EAAF are regularly counted or surveyed, for example, many of the sites covered by the AWC, the China Coastal Waterbird Census and in the lower Yangtze basin in the PRC, meaning that relatively comprehensive data was available for some sites to be assessed using our RFI methodology. Few wetland sites in Asia are intensively monitored, for example, the Luannan-Zuidong coast in Hebei Province, Chongming Dongtan in Shanghai and Mai Po in Hong Kong, China and parts of the Inner Gulf of Thailand, but these are very much the exception. Other than a few well-known sites, many wetlands in Southeast Asia are even more sporadically monitored, and the majority of sites have been counted only occasionally, perhaps only once or twice in the past 10 years, if not less.

Regularly monitored sites tend to have higher maximum counts of waterbirds and longer lists of species that regularly exceed the relevant 1% population thresholds. The PC1 scores of many of the lesser-known sites will be perennially underestimated because of the incomplete data available on migratory waterbirds, but some of these sites will nevertheless have been selected as priority sites based on this limited data available. Sites with extensive areas of potential habitat for waterbirds, but lacking count data, should be included as priorities for survey in the future.

### Considering ecological connectivity

The Convention on Migratory Species (CMS) assessed that the conservation needs of migratory species can be best represented in the newly adopted Global Biodiversity Framework through stronger consideration for ecological connectivity through management of networks of critically important sites and habitats used by migratory birds^87^. Satellite-tracking studies of migratory shorebirds have shown that species in the region utilised wetlands not previously known as important sites (e.g., Great Knot *Calidris tenuirostris*^88^). There is a risk that little known stopover sites might be lost with deleterious effects on migratory connectivity even before they are properly surveyed and studied, given that each species uses a different suite and network of wetland sites during the breeding period, northward and southward (passage) migrations and during the non-breeding (northern wintering) period. Recent studies have investigated how the concept of ecological connectivity can be applied to migratory species (e.g., Xu *et al.*^83,89^), but the data available to assess connectivity in the East Asian-Australasian Flyway, is currently limited^36^ particularly in Southeast Asia.

Further investigations will be needed to consider whether globally threatened waterbirds are adequately represented in the prioritised RFI wetlands during the non-breeding period or if important sites for these species in the 10 RFI countries are missed. For example, the coverage afforded to the Endangered Spotted Greenshank *Tringa guttifer* by the 147 RFI priority sites (Fig. 3) has showed the (relative) irreplaceability of the wetland sites selected for that species. We will also continue to review new publications on ecological connectivity and investigate the feasibility of applying this concept.

**Figure 3.**
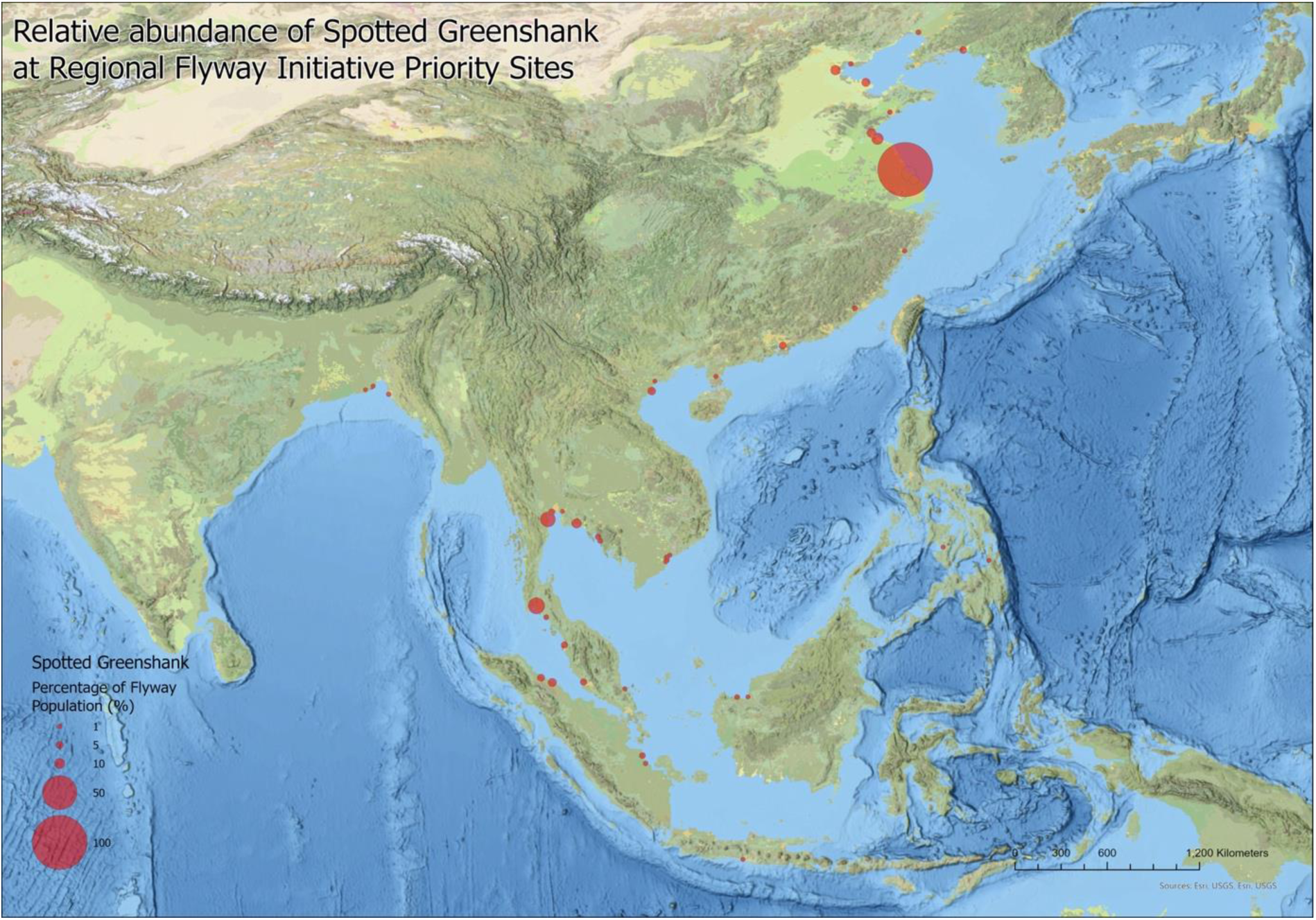
Representation of the Spotted Greenshank in Regional Flyway Initiative (RFI) priority sites across the East Asian-Australasian Flyway (EAAF).

### Implications for policy development and management

The selection of priority sites reveals a notable discrepancy between the prevalence of coastal and inland wetlands in the list. These distinct wetland types require nuanced policy development and management approaches. Coastal wetlands, abundant among the chosen sites within countries such as Indonesia and Viet Nam, possess attributes such as storm buffering capabilities that protect coastlines from erosion and tidal surges^90^. Policies must prioritise their conservation and restoration to strengthen coastal resilience. The crucial balance of salinity within these tidal wetlands also requires targeted policies to counteract threats such as industrial pollution and agricultural runoff as well as reduced flows of fresh water from rivers.

In contrast, the prominence of inland wetlands within some countries such as Cambodia underscores their role in flood regulation, absorbing excess water during heavy precipitation events^90^. Effective policies should acknowledge their importance in mitigating flood risks and restrict urban development in vulnerable regions. The provision of drinking water by inland wetlands further underscores the need for policies to curb nearby pollution sources.

A substantial number of selected priority sites already have some protection status, either as designated protected areas or with substantial overlap with such areas. Future investments in these protected sites and specific management actions that benefit requirements of migratory birds offer opportunities to enhance protection for migratory birds, support local livelihoods, and strengthen climate resilience. However, a strategic focus on unprotected sites for future investments can effectively allocate resources where management efforts are needed. Such investments may encourage the designation of these sites in future which also help governments meet the 30x30 target of the Global Biodiversity Framework.

Contextualising these and other important sites for waterbirds within the flyway and visualising them could be undertaken through expansion of the Critical Site Network Tool^91,92^, developed by Wetlands International and BirdLife International, to the EAAF.

Considering the socioeconomic context, many priority sites are in countries classified as having very high or high human development tiers. Therefore, this presents an avenue for strategic investment targeting sites in Bangladesh, Cambodia, the Lao PDR, and the Philippines, four countries in the medium human development tier. Such an approach would ensure the preservation of ecosystem services provided by these wetlands, consequently contributing to an improved overall standard of living for their beneficiaries. Balancing these considerations will be essential as the participating countries navigate the policy development and management and restoration for their vital wetlands.

## Conclusions

The site-based network approach, as explored in this study, contributes to the larger effort of biodiversity conservation. While not a panacea, when executed at a meaningful scale, it has the potential to stabilise and restore populations of some migratory bird species. The RFI underscores the need for a multipronged conservation strategy that encompasses species-focused interventions, site-based efforts, and policy-driven approaches. By embracing this holistic perspective, we aspire to catalyse lasting positive change in wetland conservation in the EAAF.

We have developed quantitative criteria for choosing priority sites for RFI in 10 participating countries. Going forward, there is a need to ensure that the portfolios of wetland sites and landscapes are congruent with the priorities of each country in relation to national conservation strategies, development plans and various international obligations and resourcing opportunities. There is also a need to assess priority sites for their importance to various ecosystem services, in order to build a strong case for long-term management, including initiatives that will result from the implementation of RFI. The next steps include (1) further consultation with government and civil society stakeholders to prioritise sites (for potential project development), delineate and agree on site boundaries, if needed; (2) assessment of ecosystem services provided by the priority sites; and (3) dissemination of RFI prioritisation findings to relevant stakeholders, and hence enhance ecological connectivity of the priority sites^93^. Through this comprehensive approach, the RFI seeks to safeguard the dynamic interplay between migratory waterbirds, wetlands ecosystems, and local communities within the EAAF.

## Supplementary Information

Supplementary Table 1

Supplementary Table 2

Supplementary Table 3

## Acknowledgements

We are greatly indebted to colleagues, friends, and partners who have contributed to and have been consulted in the conduct of the site selection process. We thank the national coordinators and many volunteer networks of the Asian Waterbird Census who have provided the annual counts to Wetlands International and other national and regional experts for providing data on waterbirds and wetland sites in specific countries: Nyambayar Batbayar, Bou Vorsak, Bui Thanh Trung, Annadel Cabanban, Hong Chamnan, Jimmy Choi, Gregorio de la Rosa, Krairat Eiamampai, Ragil Satriyo Gumilang, Amarkhuu Gungaa, Enam Ul Haque, Ferry Hasudungan, Ayuwat Jearwattanakanok, Le Trong Trai, Mike Lu, Samphors Ly, David Melville, Muntjargal Myagmar, Yus Rusila Noor, Nguyen Hoai Bao, Nguyen Quang Hao, Nguyen Van Thang, Phan Van Truong, Lisa M. Paguntalan, Gankhuyag Purev-Ochir, Chairunas Adha Putra, Josiah David Quimpo, Philip Round, Jacelyn See, Khwankhao Sinhaseni, Gombobaatar Sundev, Anson Tagtag, Batrisyia Teepol, Niyom Thongmuean, Sonny Wong, Santi Xayyasith, Xia Shaoxia, Yeap Chin Aik, Yu Xiubo, and Yu Yat-tung.

The views expressed in this publication are those of the authors and do not necessarily reflect the views and policies of the Asian Development Bank (ADB) or its Board of Governors or the governments they represent. ADB does not guarantee the accuracy of the data included in this publication and accepts no responsibility for any consequence of their use. The mention of specific companies or products of manufacturers does not imply that they are endorsed or recommended by ADB in preference to others of a similar nature that are not mentioned. By making any designation of or reference to a particular territory or geographic area in this document, ADB does not intend to make any judgments as to the legal or other status of any territory or area.

## Author Contributions

Conceptualisation: MC, DL

Data curation: MC, SW, DLY, GA, SC, GX, WH, AJ, CL, YL, TM, HO, SJ, TT, DW, ZQ, LB, KO, KP

Formal analysis: MC, SW, DLY, KP

Funding acquisition: DL

Investigation: All authors

Methodology: All authors

Writing – original draft: MC, SW, DLY, KP

Writing – review & editing: All authors

