## Supplementary material for "An innovative approach in identifying a network of priority wetland sites in the East Asian-Australasian Flyway for a new sustainable management investment programme": Crosby et al - SUPPLEMENTARY INFORMATION.docx

**Supplementary Table 1.** RFI waterbird count data file structure

| **Field name** | **Purpose of field** |
| --- | --- |
| Seq | Code from 1 to X, to allow file to be reordered in original taxonomic sequence |
| Reference | Citation of the source of waterbird count data |
| English name | Waterbird species name |
| Scientific name | Waterbird species name |
| IUCN | IUCN Red List category (CR; EN; VU; NT; LC) |
| Country: Province | Country name, plus province in large countries |
| Locality name | Name of recording locality |
| Site name | Name of the site (may include several localities) |
| Count | Waterbird count (maximum or average maximum count) |
| Selected count | The count of a species at a site selected to calculate the CSR1 Score |
| Date | Date(s) of waterbird count |
| Primary Source | Primary source of waterbird count data, if given in the Reference |
| CSR1 | Conservation Status Review (2022) 1% threshold |
| CSR1 Score | Species score using CSR1 1% population threshold |

**Supplementary Table 2.** Sources of waterbird count and site data for the 10 participating countries*.*

| **Country** | **Number of coastal and/or inland sites assessed** | **Sources of count and site data** |
| --- | --- | --- |
| Bangladesh | >8* | **Asian Waterbird Census (AWC), project reports, peer-reviewed papers.**  Data from the Asian Waterbird Census dataset was provided by Enam Ul Haque (AWC National Coordinator) and Samiul Mohsanin, which formed the basis of the information collated for site prioritisation. More than 70 sites were covered in the annual AWC for Bangladesh, including many localities counted at relatively fine spatial scales. In consultation with Sayam Chowdhury (Bangladesh Spoon-billed Sandpiper project lead) and Enam Ul Haque, we were able to reduce the number of counting localities included in the analysis. Additional data was compiled from Spoon-billed Sandpiper (CR) surveys and peer-reviewed articles, including material reviewing the status of threatened species in the country such as Masked Finfoot (CR) (Chowdhury et al. 2020).  * The AWC data for Bangladesh covered 178 individual counting localities (mostly with only low numbers of waterbirds), but (in consultation with national experts) we did not define sites to include these, other than for the eight Priority Sites. |
| Cambodia | 12 | **Asian Waterbird Census, project reports from Wildlife Conservation Society (WCS) and BirdLife International, peer-reviewed papers**  Data from the Asian Waterbird Census dataset formed the bulk of the information collated for site prioritisation, and was supplemented by published project reports and a few peer-reviewed papers. About 12 sites are covered in the annual AWC for Cambodia, although this tends to focus on wetland sites around the floodplains of the Tonle Sap, and the three main wetlands known for Sarus Crane. Hong Chamnan and Bou Vorsak, the co-coordinators of the AWC, facilitated the sharing of the AWC count data for 2020-2021 for Cambodia, including targeted counts of waterbirds at sites such as Tonle Sap and Boueng Chhmar. We were also able to access reports and the conservation strategy for the Southeast Asian subspecies of Sarus Crane (*Grus antigone sharpii*), which provides high-resolution counts of cranes over the past decade, as well as Sarus Crane census reports jointly produced by WCS, International Crane Foundation and BirdLife Cambodia. Two unpublished reports by NatureLife Cambodia on the status of shorebirds based on joint surveys with BirdLife International provided useful recent data for the assessment of the Koh Kapik Ramsar Site, possibly the most important coastal wetland in Cambodia, although some poorly known sites on the south-east coast (i.e., Kaeb, Kampot provinces) are currently being surveyed under a separate project. |
| PRC (coastal) | 66 | **China Coastal Waterbird Census, Yellow-Sea Bohai Coordinated Waterbird Surveys by Wetlands International, Annual Black-faced Spoonbill Census, EAAFP Site Information Sheets, BirdLife Datazone, Paulson Institute *Blueprint of Coastal Wetland Conservation and Management in China* (中国滨海湿地保护管理战略研究), project reports, peer-reviewed papers**  Important data sets evaluated for the PRC included the China Coastal Waterbird Census (CCWC), a large-scale monitoring initiative spearheaded by the BirdLife partner, the Hong Kong Bird Watching Society in collaboration with other Chinese birdwatching societies, and count data (and associated reports) by Wetlands International China and the Paulson Institute’s blueprint of coastal wetlands blueprint. Additional data was compiled from the peer-reviewed literature, including several studies such as Xia et al. (2017) and Duan et al. (2020), which have conducted analyses to prioritise wetland sites on the PRC coast. Species-specific studies published from 2010 onwards, including census reports (for Black-faced Spoonbill and Spoon-billed Sandpiper), also held considerable raw count data at the site level, which were factored into the calculations of scores for each site. |
| PRC (inland) | 33 | **Peer-reviewed papers, Reports on the coordinated surveys for wintering waterbirds of Central and Lower Yangtze, EAAFP Site Information Sheets, BirdLife Datazone**  For inland wetlands, the articles published on waterbird populations in a special edition of Wildfowl in 2020 were a key source for sites in the Yangtze basin, and peer-reviewed articles were the main source for inland sites in the Yellow River basin and northeast PRC. Additional input on the importance of specific sites was provided by five bird experts, namely Shi Jianbin, Gan Xiaojing, Jimmy Choi, Terry Townshend and Yu Yat-tung. |
| Indonesia | 24 | **Asian Waterbird Census, project reports from EAAFP and Manfred Stiftung/Eksai foundations, peer-reviewed papers**  Data from peer-reviewed studies, the AWC and EAAFP project reports formed the bulk of the data evaluated for Indonesia. More than 50 papers and associated documents (including verified eBird lists) were compiled, and raw count data for species at the site level were extracted for these studies. BirdLife International has supported several surveys of the eastern Sumatran coast from 2019 to 2022 in the provinces of Aceh and North Sumatra, and the data collected from this field work was very useful for several poorly surveyed coastal sites. Four bird experts, Chairunas Adha Putra, Yus Rusila Noor, Ferry Hasudungan, and Ragil Satriyo Gumilang, provided additional feedback and species data on specific sites. |
| Lao PDR | 5 | **Project reports, IUCN and Wildlife Conservation Society (WCS) reports, BirdLife/WCS survey datasets**  Limited information was available for the assessment of Laotian wetland sites, and several data sources are very dated. A key source of data for our assessment of Laotian wetlands is the ASEAN Flyway Network project, which carried out comprehensive surveys of wetlands along the Vientiane stretch of the Mekong in 2018-2020. In addition, we were able to access several project reports, kindly shared with us by one bird expert, Santi Xayyasith. Additional sources of data included biodiversity project reports on the Xe Pian-Dong Kanthung region, and peer-reviewed papers. |
| Malaysia | 9 | **Asian Waterbird Census, peer-reviewed articles, ASEAN Flyway project survey reports**  Given its wide coverage of count sites in Malaysia’s different states, the Asian Waterbird Census formed the single most important source of count data for the assessment of Malaysian wetland sites. The data are managed by the Malaysian Nature Society, the BirdLife International Partner, and for the purpose of this assessment, datasets from recent years from 2015 to 2021 were released for analyses. Additional data was obtained from project reports and peer-reviewed papers from ongoing shorebird projects in Penang, Sabah, and Sarawak. Input on species counts and sites was provided by three waterbird experts, namely Yeap Chin Aik, Jacelyn See, and Batrisyia Teepol. |
| Mongolia | 48 | **Peer-reviewed papers, IBA datasheets and Mongolian Red Data Book (birds), expert reviews**  There was relatively little recent count data available for most sites in Mongolia. To address this knowledge gap, we calculated scores for each relevant wetland IBA for Mongolia based on species identified to have exceeded their 1% thresholds in the Mongolian inventory for IBAs (Batbayar & Tseveenmyadag 2009), and site information sheets of the EAAFP Flyway Network sites. This formed the basis of our prioritisation work in Mongolia. In addition, we supplemented the Mongolian data with raw species counts from peer-reviewed literature on waterbirds in selected sites in the country, such as Ganga Lakes, Khukh Lake and the Darkhad Depression. Additional input on the ranking of sites in the preliminary Mongolia site list were provided by key national ornithological experts, including Nyambayar Batbayar, Sundev Gombobaatar, Gankhuyag Purev-Ochir and Amarkhuu Gungaa. |
| Philippines | 20 | **Asian Waterbird Census, ASEAN Flyway Network project reports, peer-reviewed papers, project reports**  The annual Asian Waterbird Census formed the most important source of data for the Philippine site assessment, and we were able to assess raw count data collated for the whole country through the approval of the AWC coordinators, Anson Tagtag (DENR-BMB), Annabel Cabanban (Wetlands International Philippines Programme) and Mike Lu (Wild Bird Club of the Philippines). Additional data were obtained from ASEAN Flyway Project reports, verified eBird lists, and peer-reviewed papers. Arne Jensen, a coordinator of the Asian Waterbird Census and a consultant for Wetlands International Philippines was responsible for compiling the Philippine AWC dataset from 2018 to 2021, and prepared guidance on the use and analyses of counts and site definitions. Cynthia Layusa provided additional information on seabird colonies on offshore islets. |
| Thailand | 18 | **Asian Waterbird Census, Bird Conservation Society of Thailand, project reports, EAAFP Site Information Sheets, BirdLife Datazone**  The Asian Waterbird Census formed the largest source of data for analyses for Thailand. We were able to access the AWC summaries and count data through the permission and coordination of the Bird Conservation Society of Thailand (BCST) and the Department of Wildlife and National Parks (DNP), the lead government agency overseeing the coordination of the AWC in Thailand. Reports from ongoing work by BCST, including a nation-wide survey of Spotted Greenshank and annual Spoon-billed Sandpiper census reports were also used for the compilation of raw count data. Philip Round and Khwankhao Sinhaseni provided input to developing a preliminary list of key sites for the assessment. |
| Viet Nam | 14 | **Black-faced Spoonbill annual census, Asian Waterbird Census, Spoon-billed Sandpiper annual census, Viet Nature and Mekong shorebird project reports, Spoon-billed Sandpiper Task Force tracking data**  Very little data from the Asian Waterbird Census (AWC) exists for Viet Nam, and only a handful of sites are regularly counted, all in the Red River Delta. Xuan Thuy National Park is one of the sites regularly counted, led by Viet Nature. However, our datasets for the analyses benefited from ongoing BirdLife and Viet Nature projects carried out in both the Red River Delta and the Mekong Delta, which provided recent count data. Additional data was compiled from project reports covering sites such as Thai Thuy and Tram Chim, while we were also able to obtain data from annual censuses of Sarus Crane, Spoon-billed Sandpiper, and Black-faced Spoonbill *Platalea minor*. Three experts provided useful feedback on sites and species, namely Le Trong Trai, Nguyen Hoai Bao, and Nguyen Quang Hao. |
| Total | >257 |  |

**Supplementary Table 3.** Information on priority sites in the 10 participating countries, including the site name and province; central coordinates; species that exceed 1% of the EAAF population at the site; IUCN Red List status for globally threatened species (CR = Critically Endangered; EN = Endangered; VU = Vulnerable); outstanding waterbird population (* = exceeding 10% of CSR1 estimate; ** = exceeding 50% of CSR1 estimate); Ramsar site designation (^R^); Flyway Network Site designation (^F^); (RFI priority site score (see Material and methods section for explanation of scoring system).

***Bangladesh***

| **SITE** | **EAAF SPECIES (≥1%)** | **SCORE** |
| --- | --- | --- |
| **Eastern Sundarbans** ^R^, Khulna  **(**[**22.00 N, 89.70 E**](https://goo.gl/maps/Bci58Zo79zuBxm6J6)**)** | Ruddy Shelduck *Tadorna ferruginea*  Masked Finfoot *Heliopais personatus** (CR)  Lesser Sandplover *Charadrius mongolus*^1^ | 42.0 |
| **Eastern Meghna Delta**, Chittagong  **(**[**22.47 N, 91.27 E**](https://goo.gl/maps/2URr4ypGLwBBiWMT7)**)** | Indian Skimmer *Rynchops albicollis** (EN)  Lesser Sandplover *Charadrius mongolus*  Spoon-billed Sandpiper *Calidris pygmaea* (CR)  Broad-billed Sandpiper *Calidris falcinellus* | 30.3 |
| **Hatia Island (including Nijhum Dwip National Park)**, Chittagong  **(**[**22.08 N, 91.07 E**](https://goo.gl/maps/E5Uxr7vYCsFhy3R7A)**)** | Indian Skimmer *Rynchops albicollis** (EN)  Lesser Sandplover *Charadrius mongolus*  Black-headed Ibis *Threskiornis melanocephalus* | 16.9 |
| **Tanguar Haor and Panabeel** ^RF^, Sylhet  **(**[**25.13 N, 91.03 E**](https://goo.gl/maps/8Q2EnSiBfZauyi8Q8)**)** | Ferruginous Duck *Aythya nyroca*  Garganey *Spatula querquedula*  Black-tailed Godwit *Limosa limosa*  Gadwall *Mareca strepera*  Common Teal *Anas crecca* | 16.2 |
| **Hakaluki Haor** ^F^, Sylhet  **(**[**24.65 N, 92.08 E**](https://goo.gl/maps/FoGL7zP456JPCtQB7)**)** | Fulvous Whistling-duck *Dendrocygna bicolor*  Ferruginous Duck *Aythya nyroca*  Black-headed Ibis *Threskiornis melanocephalus*  Asian Openbill *Anastomus oscitans* | 7.6 |
| **Sonadia Island** ^F^, Chittagong  **(**[**21.50 N, 91.88 E**](https://goo.gl/maps/6B6DgLM32RJYn5dv9)**)** | Lesser Sandplover *Charadrius mongolus*  Spoon-billed Sandpiper *Calidris pygmaea* (CR) | 4.9 |
| **Central Meghna Delta**, Barisal  **(**[**21.93 N, 90.62 E**](https://goo.gl/maps/EVQzwWnfbzoqAaST9)**)** | Black-headed Ibis *Threskiornis melanocephalus*  Lesser Sandplover *Charadrius mongolus*  Common Shelduck *Tadorna tadorna* | 3.5 |
| **Hail Haor (including Bakka Beel)** ^F^, Sylhet  **(**[**24.37 N, 91.68 E**](https://goo.gl/maps/2icovnEz53HHRt7AA)**)** | Pacific Golden Plover *Pluvialis fulva* | 1.1 |

^1^ Nearly meeting 1% of CSR1 estimate

^*^ Exceeding 10% of CSR1 estimate

Sites in **bold** overlap with a protected area(s)

***Cambodia***

| **SITE** | **EAAF SPECIES (≥1%)** | **SCORE** |
| --- | --- | --- |
| **Prek Toal (part of Tonle Sap Biosphere Reserve)** ^R^, Battambang  **(**[**13.12 N, 103.65 E**](https://goo.gl/maps/QaipWZRKoDLKbV1L7)**)** | Spot-billed Pelican *Pelecanus philippensis* **  Painted Stork *Mycteria leucocephala* **  Greater Adjutant *Leptoptilos dubius* (EN) **  Asian Openbill *Anastomus oscitans*  Masked Finfoot *Heliopais personatus* (CR) (estimates only) | 250.7 |
| **Ang Tropeang Thmor (Sarus Crane Reserve)**, Siem Reap  **(**[**13.82 N, 103.31 E**](https://goo.gl/maps/9xhjo1yXVZH2VEZT9)**)** | Sarus Crane *Grus antigone sharpii* (VU)*  Garganey *Spatula querquedula*  Spot-billed Pelican *Pelecanus philippensis*  Painted Stork *Mycteria leucocephala*  Black-headed Ibis *Threskiornis melanocephalus* | 33.3 |
| **Boeung Prek Lapouv (Sarus Crane Reserve)**, Takeo  **(**[**10.72 N, 105.03 E**](https://goo.gl/maps/PNTQDzR5tEhDToxq9)**)** | Sarus Crane *Grus antigone sharpii* (VU)*  Painted Stork *Mycteria leucocephala* | 30.4 |
| **Anlung Pring (Sarus Crane Reserve)** ^F^, Kampot  **(**[**10.48 N, 104.53 E**](https://goo.gl/maps/TMJj3dC8JheSdzEd6)**)** | Sarus Crane *Grus antigone sharpii* (VU)*  Black-tailed Godwit *Limosa limosa* | 21.3 |
| **Boeng Chhmar Ramsar Site** ^R^, Kampong Thom  **(**[**12.83 N, 104.29 E**](https://goo.gl/maps/ATE1QzUdGuX8cRhz7)**)** | Painted Stork *Mycteria leucocephala* *  Greater Adjutant *Leptoptilos dubius* (EN)  Spot-billed Pelican *Pelecanus philippensis* | 19.4 |
| **Chi Kreng**, Siem Reap  **([13.01 N, 104.42 E](https://goo.gl/maps/uZqSttuSFZHJ2UkU9))** | Painted Stork *Mycteria leucocephala*  Sarus Crane *Grus antigone sharpii* (VU) | 5.4 |
| **Kulen Promtep Wildlife Sanctuary (Memey River)**, Preah Vihear  **(**[**14.02 N, 104.52 E**](https://goo.gl/maps/DVeQsMqJ4t6YvFm18)**)** | Masked Finfoot *Heliopais personatus* (CR) | 4.7 |
| **Koh Kapik (Peam Krasop Wildlife Sanctuary)** ^R^, Koh Kong  **(**[**11.54 N, 102.97 E**](https://goo.gl/maps/DXqDWhYBWmVr16tv8)**)** | Lesser Sandplover *Charadrius mongolus*  Spotted Greenshank *Tringa guttifer* (EN) | 3.2 |
| Stoung, Kampong Thom  **(**[**12.99 N, 104.47 E**](https://goo.gl/maps/ghEZxer2yv7d32Kb9)**)** | Sarus Crane *Grus antigone sharpii* (VU) | 1.0 |

* Exceeding 10% of CSR1 estimate

** Exceeding 50% of CSR1 estimate

Sites in **bold** overlap with a protected area(s)

***Coastal People’s Republic of China***

**Liaoning Province**

| **SITE** | **EAAF SPECIES (≥1%)** | **SCORE** |
| --- | --- | --- |
| **Yalu Jiang estuary ^F^**  **([39.83 N, 124.10 E](https://goo.gl/maps/YHMak4dJJXcQdLa76))** | Swan Goose *Anser cygnoid* (VU)  Greater White-fronted Goose *Anser albifrons*  Goosander *Mergus merganser*  Common Shelduck *Tadorna tadorna*  Mandarin Duck *Aix galericulata*  Tufted Duck *Aythya fuligula*  Hooded Crane *Grus monacha* (VU)*  Chinese Egret *Egretta eulophotes* (VU)  Eurasian Oystercatcher *Haematopus ostralegus**  Grey Plover *Pluvialis squatarola*  Kentish Plover *Charadrius alexandrinus* *  Lesser Sandplover *Charadrius mongolus*  Whimbrel *Numenius phaeopus*  Eurasian Curlew *Numenius arquata*  Far Eastern Curlew *Numenius madagascariensis* (EN) *  Bar-tailed Godwit *Limosa lapponica* **  Black-tailed Godwit *Limosa limosa*  Ruddy Turnstone *Arenaria interpres*  Great Knot (EN) *Calidris tenuirostris* *  Broad-billed Sandpiper *Calidris falcinellus*  Sharp-tailed Sandpiper *Calidris acuminata* (VU)  Curlew Sandpiper *Calidris ferruginea* *  Spoon-billed Sandpiper *Calidris pygmaea* (CR)  Dunlin *Calidris alpina*  Terek Sandpiper *Xenus cinereus*  Spotted Redshank *Tringa erythropus*  Common Greenshank *Tringa nebularia*  Spotted Greenshank *Tringa guttifer* (EN)  Saunders's Gull *Saundersilarus saundersi* (VU)  Relict Gull *Larus relictus* (VU)*  Mew Gull *Larus canus* | 308.66 |
| **Liaohe Estuary National Nature Reserve and Inner Gulf of Liaodong ^RF^**  **(**[**40.90 N, 121.78 E**](https://goo.gl/maps/dhbdYRKdhfF66gEx5)**)** | Bean Goose *Anser fabalis*  Common Shelduck *Tadorna tadorna*  Falcated Duck *Mareca falcata*  Siberian Crane *Leucogeranus leucogeranus* (CR)  Red-crowned Crane *Grus japonensis* (VU)**  Hooded Crane *Grus monacha* (VU)*  Oriental Stork *Ciconia boyciana* (EN)  Eurasian Spoonbill *Platalea leucorodia*  Great Cormorant *Phalacrocorax carbo*  Eurasian Oystercatcher *Haematopus ostralegus**  Black-winged Stilt *Himantopus himantopus*  Grey Plover *Pluvialis squatarola*  Kentish Plover *Charadrius alexandrinus* *  Lesser Sandplover *Charadrius mongolus* *  Eurasian Curlew *Numenius arquata*  Far Eastern Curlew (EN) *Numenius madagascariensis*  Ruddy Turnstone *Arenaria interpres*  Great Knot (EN) *Calidris tenuirostris* *  Red Knot *Calidris canutus* *  Sharp-tailed Sandpiper *Calidris acuminata* (VU)  Dunlin *Calidris alpina*  Terek Sandpiper *Xenus cinereus*  Spotted Redshank *Tringa erythropus **  Common Greenshank *Tringa nebularia*  Common Redshank *Tringa totanus*  Marsh Sandpiper *Tringa stagnatilis*  Spotted Greenshank *Tringa guttifer* (EN)  Saunders's Gull *Saundersilarus saundersi* (VU) *  Relict Gull *Larus relictus* (VU)  Mew Gull *Larus canus*  Little Tern *Sternula albifrons* | 272.71 |
| **Zhuanghe coast**  **(**[**39.67 N, 123.03 E**](https://goo.gl/maps/e9LcJvXKgVrdzP9a9)**)** | Black-faced Spoonbill *Platalea minor* (EN)  Chinese Egret *Egretta eulophotes* (VU)  Whimbrel *Numenius phaeopus*  Little Curlew *Numenius minutus*  Eurasian Curlew *Numenius arquata*  Far Eastern Curlew *Numenius madagascariensis* (EN)  Bar-tailed Godwit *Limosa lapponica*  Black-tailed Godwit *Limosa limosa*  Relict Gull *Larus relictus* (VU)  Black-tailed Gull *Larus crassirostris*  Mew Gull *Larus canus* | 30.44 |
| **Pulandian-Jinzhou east coast, including Changshan Islands**  **(**[**39.37 N, 122.30 E**](https://goo.gl/maps/pStyPACgmtawhLkd8)**)** | Whimbrel *Numenius phaeopus*  Eurasian Curlew *Numenius arquata*  Far Eastern Curlew *Numenius madagascariensis* (EN)  Bar-tailed Godwit *Limosa lapponica*  Great Knot (EN) *Calidris tenuirostris*  Dunlin *Calidris alpina* | 27.0 |
| **Jinzhou Bay, Dalian**  **(**[**39.17 N, 121.60 E**](https://goo.gl/maps/49WaSpxNhDzUoAMd7)**)** | Bar-tailed Godwit *Limosa lapponica*  Spotted Redshank *Tringa erythropus* | 19.80 |

**Hebei Province**

| **SITE** | **EAAF SPECIES (≥1%)** | **SCORE** |
| --- | --- | --- |
| **Luannan-Zuidong coast**  **(**[**39.10 N, 118.20 E**](https://goo.gl/maps/owLxzFz3LHvSmtnn9)**)** | Common Shelduck *Tadorna tadorna*  Pied Avocet *Recurvirostra avosetta* *  Black-winged Stilt *Himantopus himantopus*  Grey Plover *Pluvialis squatarola*  Kentish Plover *Charadrius alexandrinus*  Eurasian Curlew *Numenius arquata*  Far Eastern Curlew *Numenius madagascariensis* (EN)  Black-tailed Godwit *Limosa limosa* *  Great Knot (EN) *Calidris tenuirostris*  Red Knot *Calidris canutus* **  Broad-billed Sandpiper *Calidris falcinellus*  Sharp-tailed Sandpiper *Calidris acuminata* (VU)  Curlew Sandpiper *Calidris ferruginea*  Red-necked Stint *Calidris ruficollis*  Sanderling *Calidris alba* *  Dunlin *Calidris alpina*  Asian Dowitcher *Limnodromus semipalmatus*  Spotted Redshank *Tringa erythropus*  Marsh Sandpiper *Tringa stagnatilis*  Spotted Greenshank *Tringa guttifer* (EN)  Relict Gull *Larus relictus* (VU)  White-winged Tern *Chlidonias leucopterus* | 199.05 |
| **Huanghua-Cangzhou coast**  **(**[**38.43 N, 117.67 E**](https://goo.gl/maps/8DAYgjj6V2jnHYiE7)**)** | Greylag Goose *Anser anser*  Bean Goose *Anser fabalis*  Smew *Mergellus albellus*  Common Shelduck *Tadorna tadorna*  Ruddy Shelduck *Tadorna ferruginea*  Baer's Pochard *Aythya baeri* (CR)  Falcated Duck *Mareca falcata*  Great Crested Grebe *Podiceps cristatus*  White-naped Crane *Grus vipio* (VU)  Common Crane *Grus grus*  Oriental Stork *Ciconia boyciana* (EN)  Dalmatian Pelican *Pelecanus crispus*  Eurasian Oystercatcher *Haematopus ostralegus*  Pied Avocet *Recurvirostra avosetta*  Black-winged Stilt *Himantopus himantopus*  Grey Plover *Pluvialis squatarola*  Little Ringed Plover *Charadrius dubius*  Kentish Plover *Charadrius alexandrinus*  Whimbrel *Numenius phaeopus*  Eurasian Curlew *Numenius arquata*  Far Eastern Curlew *Numenius madagascariensis* (EN)  Bar-tailed Godwit *Limosa lapponica*  Black-tailed Godwit *Limosa limosa*  Red Knot *Calidris canutus*  Sharp-tailed Sandpiper *Calidris acuminata* (VU)  Curlew Sandpiper *Calidris ferruginea*  Spotted Redshank *Tringa erythropus*  Marsh Sandpiper *Tringa stagnatilis*  Relict Gull *Larus relictus* (VU)  Mew Gull *Larus canus* | 84.61 |
| **Laoting-Caofeidian coast**  **(**[**39.02 N, 118.73 E**](https://goo.gl/maps/uiBf3M8sf9rVWsc78)**)** | Smew *Mergellus albellus*  Pied Avocet *Recurvirostra avosetta*  Kentish Plover *Charadrius alexandrinus*  Eurasian Curlew *Numenius arquata*  Far Eastern Curlew *Numenius madagascariensis* (EN)  Black-tailed Godwit *Limosa limosa*  Dunlin *Calidris alpina*  Marsh Sandpiper *Tringa stagnatilis* | 17.08 |

**Tianjin Municipality**

| **SITE** | **EAAF SPECIES (≥1%)** | **SCORE** |
| --- | --- | --- |
| **Tianjin coastal mudflats**  **(**[**38.92 N, 117.75 E**](https://goo.gl/maps/asDGNCJ2rZJRyHkb9)**)** | Whooper Swan *Cygnus cygnus*  Tundra Swan *Cygnus columbianus* *  Greylag Goose *Anser anser* *  Bean Goose *Anser fabalis*  Smew *Mergellus albellus*  Goosander *Mergus merganser*  Common Shelduck *Tadorna tadorna*  Common Pochard *Aythya ferina* (VU)  Northern Shoveler *Spatula clypeata*  Falcated Duck *Mareca falcata*  Oriental Stork *Ciconia boyciana* (EN)  Pied Avocet *Recurvirostra avosetta* *  Black-winged Stilt *Himantopus himantopus*  Grey Plover *Pluvialis squatarola*  Little Ringed Plover *Charadrius dubius*  Kentish Plover *Charadrius alexandrinus*  Lesser Sandplover *Charadrius mongolus*  Eurasian Curlew *Numenius arquata*  Far Eastern Curlew *Numenius madagascariensis* (EN)  Bar-tailed Godwit *Limosa lapponica*  Black-tailed Godwit *Limosa limosa*  Great Knot *Calidris tenuirostris* (EN)  Red Knot *Calidris canutus*  Curlew Sandpiper *Calidris ferruginea*  Red-necked Stint *Calidris ruficollis*  Dunlin *Calidris alpina*  Asian Dowitcher *Limnodromus semipalmatus*  Spotted Redshank *Tringa erythropus*  Common Greenshank *Tringa nebularia*  Marsh Sandpiper *Tringa stagnatilis*  Saunders's Gull *Saundersilarus saundersi* (VU)  Relict Gull *Larus relictus* (VU)**  Little Tern *Sternula albifrons*  Caspian Tern *Hydroprogne caspia* | 174.14 |
| **Beidagang Wetland Nature Reserve ^R^**  **([38.75 N, 117.38 E](https://goo.gl/maps/hp63ntbZkonEHzJQA))** | Tundra Swan *Cygnus columbianus*  Greylag Goose *Anser anser*  Common Shelduck *Tadorna tadorna*  Baer’s Pochard *Aythya baeri* (CR)  Falcated Duck *Mareca falcata*  Gadwall *Mareca strepera*  Dalmatian Pelican *Pelecanus crispus* *  Pied Avocet *Recurvirostra avosetta*  Black-winged Stilt *Himantopus himantopus*  Grey Plover *Pluvialis squatarola*  Little Ringed Plover *Charadrius dubius*  Kentish Plover *Charadrius alexandrinus*  Eurasian Curlew *Numenius arquata*  Far Eastern Curlew *Numenius madagascariensis* (EN) | 81.19 |

**Shandong Province**

| **SITE** | **EAAF SPECIES (≥1%)** | **SCORE** |
| --- | --- | --- |
| **Yellow River Delta National Nature Reserve ^RF^**  **([37.97 N, 118.97 E](https://goo.gl/maps/wuA6oK66VBngJqf19))** | Mute Swan *Cygnus olor*  Whooper Swan *Cygnus cygnus*  Tundra Swan *Cygnus columbianus*  Greylag Goose *Anser anser*  Swan Goose *Anser cygnoid* (VU)  Bean Goose *Anser fabalis*  Smew *Mergellus albellus*  Goosander *Mergus merganser*  Common Shelduck *Tadorna tadorna*  Ruddy Shelduck *Tadorna ferruginea*  Mandarin Duck *Aix galericulata*  Common Pochard *Aythya ferina* (VU)  Baer’s Pochard *Aythya baeri* (CR)  Falcated Duck *Mareca falcata*  Gadwall *Mareca strepera*  Chinese Spot-billed Duck *Anas zonorhyncha*  Northern Pintail *Anas acuta*  Great Crested Grebe *Podiceps cristatus*  Siberian Crane *Leucogeranus leucogeranus* (CR)*  White-naped Crane *Grus vipio* (VU)*  Red-crowned Crane *Grus japonensis* (VU)*  Common Crane *Grus grus* **  Hooded Crane *Grus monacha* (VU)*  Black Stork *Ciconia nigra*  Oriental Stork *Ciconia boyciana* (EN)  Eurasian Spoonbill *Platalea leucorodia*  Dalmatian Pelican *Pelecanus crispus* **  Great Cormorant *Phalacrocorax carbo*  Eurasian Oystercatcher *Haematopus ostralegus*  Pied Avocet *Recurvirostra avosetta*  Black-winged Stilt *Himantopus himantopus*  Grey Plover *Pluvialis squatarola*  Little Ringed Plover *Charadrius dubius*  Kentish Plover *Charadrius alexandrinus*  Lesser Sandplover *Charadrius mongolus*  Whimbrel *Numenius phaeopus*  Little Curlew *Numenius minutus*  Eurasian Curlew *Numenius arquata*  Far Eastern Curlew *Numenius madagascariensis* (EN)*  Bar-tailed Godwit *Limosa lapponica*  Black-tailed Godwit *Limosa limosa* *  Great Knot *Calidris tenuirostris* (EN)  Red Knot *Calidris canutus*  Sharp-tailed Sandpiper *Calidris acuminata* (VU)  Dunlin *Calidris alpina*  Spotted Redshank *Tringa erythropus*  Common Greenshank *Tringa nebularia*  Spotted Greenshank *Tringa guttifer* (EN)  Saunders’s Gull *Saundersilarus saundersi* (VU)*  Relict Gull *Larus relictus* (VU)*  Common Gull-billed Tern *Gelochelidon nilotica*  Caspian Tern *Hydroprogne caspia*  Common Tern *Sterna hirundo* | 595.48 |
| **Qingdao coast and Jiaozhou Bay**  **(**[**36.18 N, 120.20 E**](https://goo.gl/maps/dCxnDtFNCA8CzVUZA)**)** | Common Shelduck *Tadorna tadorna*  Eurasian Oystercatcher *Haematopus ostralegus*  Pied Avocet *Recurvirostra avosetta*  Grey Plover *Pluvialis squatarola*  Kentish Plover *Charadrius alexandrinus*  Lesser Sandplover *Charadrius mongolus*  Eurasian Curlew *Numenius arquata*  Dunlin *Calidris alpina*  Spotted Redshank *Tringa erythropus*  Spotted Greenshank *Tringa guttifer* (EN)  Saunders’s Gull *Saundersilarus saundersi* (VU)  Caspian Tern *Hydroprogne caspia*  Chinese Crested Tern *Thalasseus bernsteini* (CR)* | 78.15 |
| **Laizhou Bay**  **(**[**37.10 N, 119.40 E**](https://goo.gl/maps/rKSfGp4sqB1G4cBp6)**)** | Mute Swan *Cygnus olor*  Smew *Mergellus albellus*  Goosander *Mergus merganser*  Falcated Duck *Mareca falcata*  Northern Pintail *Anas acuta*  Eurasian Curlew *Numenius arquata* | 24.16 |
| **Wudi-Zhanhua-Hekou coast (**[**38.13 N, 118.20 E**](https://goo.gl/maps/k6PPP4AdDNZj6E5e7)**)** | Kentish Plover *Charadrius alexandrinus*  Eurasian Curlew *Numenius arquata*  Far Eastern Curlew *Numenius madagascariensis* (EN)  Red Knot *Calidris canutus*  Common Greenshank *Tringa nebularia*  Relict Gull *Larus relictus* (VU) | 15.63 |
| **Rongcheng Swan Nature Reserve ^F^**  **(**[**37.25 N, 122.57 E**](https://goo.gl/maps/V5kz8U1AaeK7fZNG9)**)** | Whooper Swan *Cygnus cygnus* * | 11.85 |

**Jiangsu Province**

| **SITE** | **EAAF SPECIES (≥1%)** | **SCORE** |
| --- | --- | --- |
| Lianyungang coast  ([**34.62 N, 119.52 E**](https://goo.gl/maps/Fksr4rfckf85Se3x6)) | Common Shelduck *Tadorna tadorna*  Falcated Duck *Mareca falcata*  Red-crowned Crane *Grus japonensis* (VU)*  Dalmatian Pelican *Pelecanus crispus* **  Eurasian Oystercatcher *Haematopus ostralegus**  Pied Avocet *Recurvirostra avosetta* *  Grey Plover *Pluvialis squatarola **  Kentish Plover *Charadrius alexandrinus*  Lesser Sandplover *Charadrius mongolus **  Eurasian Curlew *Numenius arquata*  Far Eastern Curlew *Numenius madagascariensis* (EN)  Bar-tailed Godwit *Limosa lapponica*  Black-tailed Godwit *Limosa limosa* *  Great Knot *Calidris tenuirostris* (EN)  Red Knot *Calidris canutus*  Broad-billed Sandpiper *Calidris falcinellus** Sharp-tailed Sandpiper *Calidris acuminata* (VU)*  Curlew Sandpiper *Calidris ferruginea*  Long-toed Stint *Calidris subminuta*  Red-necked Stint *Calidris ruficollis*  Dunlin *Calidris alpina*  Asian Dowitcher *Limnodromus semipalmatus* **  Terek Sandpiper *Xenus cinereus*  Spotted Redshank *Tringa erythropus*  Marsh Sandpiper *Tringa stagnatilis*  Spotted Greenshank *Tringa guttifer* (EN)*  Saunders’s Gull *Saundersilarus saundersi* (VU)  Black-headed Gull *Larus ridibundus*  Relict Gull *Larus relictus* (VU)  Caspian Tern *Hydroprogne caspia* | 344.15 |
| **Tiaozini (Dongtai coast)**  **(**[**32.75 N, 120.97 E**](https://goo.gl/maps/kYe84rx1XhghuyKA7)**)** | Falcated Duck *Mareca falcata*  Black-faced Spoonbill *Platalea minor* (EN)  Dalmatian Pelican *Pelecanus crispus* **  Great Cormorant *Phalacrocorax carbo* *  Eurasian Oystercatcher *Haematopus ostralegus*  Pied Avocet *Recurvirostra avosetta*  Grey Plover *Pluvialis squatarola*  Kentish Plover *Charadrius alexandrines* *  Lesser Sandplover *Charadrius mongolus* *  Greater Sandplover *Charadrius leschenaultii*  Eurasian Curlew *Numenius arquata*  Far Eastern Curlew *Numenius madagascariensis*  (EN)  Bar-tailed Godwit *Limosa lapponica*  Black-tailed Godwit *Limosa limosa*  Great Knot *Calidris tenuirostris* (EN)  Broad-billed Sandpiper *Calidris falcinellus* Sharp-tailed Sandpiper *Calidris acuminata* (VU)  Spoon-billed Sandpiper *Calidris pygmaea* (CR)*  Red-necked Stint *Calidris ruficollis*  Sanderling *Calidris alba*  Dunlin *Calidris alpina*  Terek Sandpiper *Xenus cinereus*  Spotted Greenshank *Tringa guttifer* (EN)**  Saunders’s Gull *Saundersilarus saundersi* (VU)  Common Tern *Sterna hirundo* | 318.33 |
| **Yancheng National Nature Reserve ^RF^**  **(**[**33.72 N, 120.52 E**](https://goo.gl/maps/MTDCtHfNDMsEqZ7C7)**)** | Goosander *Mergus merganser*  Red-crowned Crane *Grus japonensis* (VU)**  Common Crane *Grus grus*  Hooded Crane *Grus monacha* (VU)  Black-faced Spoonbill *Platalea minor* (EN)  Black-crowned Night-heron *Nycticorax nycticorax*  Great Cormorant *Phalacrocorax carbo*  Eurasian Oystercatcher *Haematopus ostralegus**  Pied Avocet *Recurvirostra avosetta*  Black-winged Stilt *Himantopus himantopus*  Grey Plover *Pluvialis squatarola* *  Pacific Golden Plover *Pluvialis fulva*  Little Ringed Plover *Charadrius dubius*  Kentish Plover *Charadrius alexandrinus*  Lesser Sandplover *Charadrius mongolus*  Eurasian Curlew *Numenius arquata*  Bar-tailed Godwit *Limosa lapponica*  Black-tailed Godwit *Limosa limosa*  Ruddy Turnstone *Arenaria interpres*  Great Knot *Calidris tenuirostris* (EN)  Red Knot *Calidris canutus*  Spoon-billed Sandpiper *Calidris pygmaea* (CR)  Red-necked Stint *Calidris ruficollis*  Sanderling *Calidris alba*  Dunlin *Calidris alpina*  Terek Sandpiper *Xenus cinereus*  Spotted Redshank *Tringa erythropus*  Common Redshank *Tringa totanus*  Saunders’s Gull Saundersilarus saundersi (VU)*  Common Tern *Sterna hirundo* * | 195.70 |
| **Ganyu coast**  **(**[**34.97 N, 119.20 E**](https://goo.gl/maps/nUKMaBxqj39rMSCn8)**)** | Red-crowned Crane *Grus japonensis* (VU)  Eurasian Spoonbill *Platalea leucorodia*  Dalmatian Pelican *Pelecanus crispus*  Eurasian Oystercatcher *Haematopus ostralegus**  Pied Avocet *Recurvirostra avosetta*  Grey Plover *Pluvialis squatarola*  Kentish Plover *Charadrius alexandrinus*  Lesser Sandplover *Charadrius mongolus*  Eurasian Curlew *Numenius arquata*  Bar-tailed Godwit *Limosa lapponica*  Black-tailed Godwit *Limosa limosa*  Great Knot *Calidris tenuirostris* (EN)  Red Knot *Calidris canutus*  Broad-billed Sandpiper *Calidris falcinellus* *  Sharp-tailed Sandpiper *Calidris acuminata* (VU)*  Curlew Sandpiper *Calidris ferruginea*  Red-necked Stint *Calidris ruficollis*  Dunlin *Calidris alpina*  Asian Dowitcher *Limnodromus semipalmatus* **  Terek Sandpiper *Xenus cinereus*  Spotted Greenshank *Tringa guttifer* (EN) | 194.23 |
| **Dongling coast**  **(**[**32.15 N, 121.45 E**](https://goo.gl/maps/DGYV1en1nxnCuLGU9)**)** | Eurasian Oystercatcher *Haematopus ostralegus**  Pied Avocet *Recurvirostra avosetta*  Grey Plover *Pluvialis squatarola*  Kentish Plover *Charadrius alexandrinus*  Lesser Sandplover *Charadrius mongolus* *  Eurasian Curlew *Numenius arquata*  Far Eastern Curlew *Numenius madagascariensis*  (EN)  Bar-tailed Godwit *Limosa lapponica*  Black-tailed Godwit *Limosa limosa*  Great Knot *Calidris tenuirostris* (EN)  Red Knot *Calidris canutus*  Broad-billed Sandpiper *Calidris falcinellus* Spoon-billed Sandpiper *Calidris pygmaea* (CR)  Red-necked Stint *Calidris ruficollis*  Sanderling *Calidris alba*  Dunlin *Calidris alpina*  Terek Sandpiper *Xenus cinereus*  Spotted Greenshank *Tringa guttifer* (EN)  Saunders’s Gull *Saundersilarus saundersi* (VU)  Little Tern *Sternula albifrons* | 126.56 |
| **Rudong coast**  **(**[**32.52 N, 121.17 E**](https://goo.gl/maps/5VChnczFEXQtsaP79)**)** | Black-faced Spoonbill *Platalea minor* (EN)  Eurasian Oystercatcher *Haematopus ostralegus* Grey Plover *Pluvialis squatarola* *  Kentish Plover *Charadrius alexandrinus **  Lesser Sandplover *Charadrius mongolus* Greater Sandplover *Charadrius leschenaultii*  Whimbrel *Numenius phaeopus*  Eurasian Curlew *Numenius arquata*  Far Eastern Curlew *Numenius madagascariensis* (EN)  Bar-tailed Godwit *Limosa lapponica*  Black-tailed Godwit *Limosa limosa*  Great Knot *Calidris tenuirostris* (EN)  Red Knot *Calidris canutus*  Broad-billed Sandpiper *Calidris falcinellus* Long-toed Stint *Calidris subminuta*  Spoon-billed Sandpiper *Calidris pygmaea* (CR)  Red-necked Stint *Calidris ruficollis*  Sanderling *Calidris alba*  Dunlin *Calidris alpina*  Terek Sandpiper *Xenus cinereus*  Spotted Redshank *Tringa erythropus*  Spotted Greenshank *Tringa guttifer* (EN)*  Saunders’s Gull *Saundersilarus saundersi* (VU)  Common Tern *Sterna hirundo* | 84.02 |
| **Dongsha shoals, PRC**  **(**[**33.00 N,** **121.23E**](https://goo.gl/maps/CiFkdi7ZfSvijHbu7)**)** | Goosander *Mergus merganser*  Common Shelduck *Tadorna tadorna*  Eurasian Oystercatcher *Haematopus ostralegus* Kentish Plover *Charadrius alexandrinus*  Far Eastern Curlew *Numenius madagascariensis* (EN)  Bar-tailed Godwit *Limosa lapponica*  Red Knot *Calidris canutus**  Broad-billed Sandpiper *Calidris falcinellus* Dunlin *Calidris alpina*  Asian Dowitcher *Limnodromus semipalmatus*  Wood Sandpiper *Tringa glareola* | 41.72 |

**Shanghai Municipality**

| **SITE** | **EAAF SPECIES (≥1%)** | **SCORE** |
| --- | --- | --- |
| **Chongming Dongtan National Nature Reserve ^RF^**  **(**[**31.48 N, 122.00 E**](https://goo.gl/maps/RT3kV4aJJqLUtnNL8)**)** | Baikal Teal *Sibirionetta formosa*  Hooded Crane *Grus monacha* (VU)*  Black-faced Spoonbill *Platalea minor* (EN)  Dalmatian Pelican *Pelecanus crispus*  Pied Avocet *Recurvirostra avosetta*  Black-winged Stilt *Himantopus himantopus*  Kentish Plover *Charadrius alexandrinus*  Eurasian Curlew *Numenius arquata*  Black-tailed Godwit *Limosa limosa*  Spoon-billed Sandpiper *Calidris pygmaea* (CR)  Red-necked Stint *Calidris ruficollis*  Dunlin *Calidris alpina*  Spotted Redshank *Tringa erythropus*  Marsh Sandpiper *Tringa stagnatilis* | 43.36 |
| **Nanhui coast (including Nanhui Wetlands Park)**  **(**[**30.93 N, 121.97 E**](https://goo.gl/maps/EiMbcBK1qxXetuQQ9)**)** | Falcated Duck *Mareca falcata*  Kentish Plover *Charadrius alexandrinus*  Lesser Sandplover *Charadrius mongolus*  Long-toed Stint *Calidris subminuta*  Red-necked Stint *Calidris ruficollis*  Spotted Redshank *Tringa erythropus* | 15.72 |

**Zhejiang Province**

| **SITE** | **EAAF SPECIES (≥1%)** | **SCORE** |
| --- | --- | --- |
| **Wenzhou Bay**  **(**[**27.92 N, 120.88 E**](https://goo.gl/maps/hGYaNEvbFpC6vgP17)**)** | Common Shelduck *Tadorna tadorna*  Common Pochard *Aythya ferina* (VU)  Tufted Duck *Aythya fuligula*  Black-faced Spoonbill *Platalea minor* (EN)  Dalmatian Pelican *Pelecanus crispus* *  Great Cormorant *Phalacrocorax carbo*  Pied Avocet *Recurvirostra avosetta*  Grey Plover *Pluvialis squatarola*  Kentish Plover *Charadrius alexandrinus*  Lesser Sandplover *Charadrius mongolus*  Eurasian Curlew *Numenius arquata*  Far Eastern Curlew *Numenius madagascariensis* (EN)  Black-tailed Godwit *Limosa limosa*  Red Knot *Calidris canutus*  Broad-billed Sandpiper *Calidris falcinellus*  Sharp-tailed Sandpiper *Calidris acuminata* (VU)  Long-toed Stint *Calidris subminuta*  Spoon-billed Sandpiper *Calidris pygmaea* (CR)  Saunders’s Gull *Saundersilarus saundersi* (VU) | 94.14 |
| **Hangzhou Bay**  **(**[**30.33 N, 121.00 E**](https://goo.gl/maps/isiVrrP1mwDXAZRc8)**)** | Baer’s Pochard *Aythya baeri* (CR)  Falcated Duck *Mareca falcata*  Great Crested Grebe *Podiceps cristatus*  Dalmatian Pelican *Pelecanus crispus*  Grey Plover *Pluvialis squatarola*  Kentish Plover *Charadrius alexandrinus*  Lesser Sandplover *Charadrius mongolus*  Bar-tailed Godwit *Limosa lapponica*  Black-tailed Godwit *Limosa limosa*  Sharp-tailed Sandpiper *Calidris acuminata* (VU)  Long-toed Stint *Calidris subminuta*  Dunlin *Calidris alpina*  Spotted Redshank *Tringa erythropus*  Common Greenshank *Tringa nebularia*  Common Redshank *Tringa totanus*  Saunders's Gull *Saundersilarus saundersi* (VU) | 43.80 |

**Fujian Province**

| **SITE** | **EAAF SPECIES (≥1%)** | **SCORE** |
| --- | --- | --- |
| **Min Jiang estuary ^R^**  **([26.12 N, 119.63 E](https://goo.gl/maps/GirHaXUTcoxPam1S7))** | Swan Goose *Anser cygnoid* (VU)  Black-faced Spoonbill *Platalea minor* (EN)  Dalmatian Pelican *Pelecanus crispus* *  Pied Avocet *Recurvirostra avosetta*  Kentish Plover *Charadrius alexandrines*  Lesser Sandplover *Charadrius mongolus*  Greater Sandplover *Charadrius leschenaultii*  Eurasian Curlew *Numenius arquata*  Ruddy Turnstone *Arenaria interpres*  Spoon-billed Sandpiper *Calidris pygmaea* (CR)  Sanderling *Calidris alba*  Terek Sandpiper *Xenus cinereus*  Spotted Redshank *Tringa erythropus*  Common Tern *Sterna hirundo*  Chinese Crested Tern *Thalasseus bernsteini* (CR) | 52.36 |
| **Quanzhou Bay, including Jin Jiang estuary**  **(**[**24.85 N, 118.67 E**](https://goo.gl/maps/vAd5ZMMGKnXaWcbn7)**)** | Dalmatian Pelican *Pelecanus crispus*  Pied Avocet *Recurvirostra avosetta*  Grey Plover *Pluvialis squatarola*  Kentish Plover *Charadrius alexandrines*  Lesser Sandplover *Charadrius mongolus*  Whimbrel *Numenius phaeopus*  Eurasian Curlew *Numenius arquata*  Curlew Sandpiper *Calidris ferruginea*  Sanderling *Calidris alba*  Dunlin *Calidris alpina*  Terek Sandpiper *Xenus cinereus*  Saunders’s Gull *Saundersilarus saundersi* (VU) | 24.77 |
| **Xiamen coast**  **(**[**24.57 N, 118.08 E**](https://goo.gl/maps/jD3ck4LJxMpgTMW48)**)** | Great Cormorant *Phalacrocorax carbo*  Kentish Plover *Charadrius alexandrines*  Whimbrel *Numenius phaeopus*  Spotted Greenshank *Tringa guttifer* (EN)  Saunders’s Gull *Saundersilarus saundersi* (VU)  Caspian Tern *Hydroprogne caspia* | 13.49 |
| Xinghua Bay  **(**[**25.45 N, 119.18 E**](https://goo.gl/maps/ykR88UXMGuWy91pf8)**)** | Black-faced Spoonbill *Platalea minor* (EN)  Great Cormorant *Phalacrocorax carbo*  Kentish Plover *Charadrius alexandrines*  Dunlin *Calidris alpina*  Saunders’s Gull *Saundersilarus saundersi* (VU) | 12.94 |

**Guangdong Province**

| **SITE** | **EAAF SPECIES (≥1%)** | **SCORE** |
| --- | --- | --- |
| **Inner Deep Bay and Shenzhen River catchment ^RF^**  **(**[**22.50 N, 114.02 E**](https://goo.gl/maps/ZCSMhPdfQ4f48arB9)**)** | Tufted Duck *Aythya fuligula*  Northern Shoveler *Spatula clypeata*  Northern Pintail *Anas acuta*  Great Crested Grebe *Podiceps cristatus*  Black-faced Spoonbill *Platalea minor* (EN)  Great White Egret *Ardea alba*  Dalmatian Pelican *Pelecanus crispus*  Great Cormorant *Phalacrocorax carbo* *  Pied Avocet *Recurvirostra avosetta*  Black-winged Stilt *Himantopus himantopus*  Pacific Golden Plover *Pluvialis fulva*  Little Ringed Plover *Charadrius dubius*  Kentish Plover *Charadrius alexandrines*  Eurasian Curlew *Numenius arquata*  Black-tailed Godwit *Limosa limosa*  Curlew Sandpiper *Calidris ferruginea*  Spoon-billed Sandpiper *Calidris pygmaea* (CR)  Asian Dowitcher *Limnodromus semipalmatus*  Terek Sandpiper *Xenus cinereus*  Spotted Redshank *Tringa erythropus*  Common Greenshank *Tringa nebularia*  Common Redshank *Tringa totanus*  Marsh Sandpiper *Tringa stagnatilis*  Spotted Greenshank *Tringa guttifer* (EN) | 68.84 |
| **Haifeng wetlands ^R^**  **([22.83 N, 115.22 E](https://goo.gl/maps/eY2SLKBxRXUS5ELv9))** | Northern Pintail *Anas acuta*  Black-faced Spoonbill *Platalea minor* (EN)  Great White Egret *Ardea alba*  Dalmatian Pelican *Pelecanus crispus*  Great Cormorant *Phalacrocorax carbo*  Spotted Redshank *Tringa erythropus* | 12.78 |
| **Zhanjiang-Leizhou Peninsula coast ^R^**  **([20.85 N, 110.20 E](https://goo.gl/maps/FnpyXr2dQfzqLmKGA))** | Black-faced Spoonbill *Platalea minor* (EN)  Kentish Plover *Charadrius alexandrines*  Lesser Sandplover *Charadrius mongolus*  Broad-billed Sandpiper *Calidris falcinellus*  Spoon-billed Sandpiper *Calidris pygmaea* (CR)  Saunders’s Gull *Saundersilarus saundersi* (VU)  Caspian Tern *Hydroprogne caspia* | 12.62 |

**Hainan Province**

| **SITE** | **EAAF SPECIES (≥1%)** | **SCORE** |
| --- | --- | --- |
| **Dongzhaigang Nature Reserve ^R^**  **([20.02 N, 110.57 E](https://goo.gl/maps/gsoQ1wuSHnZrDTo97))** | Pied Avocet *Recurvirostra avosetta*  Kentish Plover *Charadrius alexandrines*  Whimbrel *Numenius phaeopus*  Sanderling *Calidris alba* | 16.69 |
| Danzhou-Lingao coast  **(**[**19.73 N, 109.23 E)**](https://goo.gl/maps/x4p32BNDdwtbVE5NA) | Lesser Sandplover *Charadrius mongolus* | 9.29 |

**Guangxi Province**

| **SITE** | **EAAF SPECIES (≥1%)** | **SCORE** |
| --- | --- | --- |
| Beihai coast ^R^  **([21.45 N, 109.32 E](https://goo.gl/maps/VvLhaQAhQeabw9id9))** | Little Ringed Plover *Charadrius dubius*  Kentish Plover *Charadrius alexandrines*  Lesser Sandplover *Charadrius mongolus*  Spoon-billed Sandpiper *Calidris pygmaea* (CR)  Sanderling *Calidris alba* | 11.01 |
| Fangcheng coast ^R^  **([21.57 N, 108.15 E](https://goo.gl/maps/tqFUbG5VE5bFdu8o6))** | Kentish Plover *Charadrius alexandrines*  Lesser Sandplover *Charadrius mongolus*  Spoon-billed Sandpiper *Calidris pygmaea* (CR) | 8.49 |

***Northeast People’s Republic of China***

**Heilongjiang Province**

| **SITE** | **IMPORTANCE OF SITE FOR MIGRATORY WATERBIRDS** |
| --- | --- |
| **Zhalong National Nature Reserve ^RF^**  **(**[**47.20 N, 124.20 E**](https://goo.gl/maps/fg1erFjpNPiPE1B76)**)** | Zhalong NNR comprises extensive reedbeds, pools and marshland that supports large numbers of breeding and migrating waterbirds, notably an important breeding population of Red-crowned Crane (VU). The wetlands there have experienced serious ecological problems and lost some of their value for migratory waterbirds, but measures are underway to reverse the trends of increasing ecological degradation (Su *et al.* 2011). |
| **Xingkai Lake National Nature Reserve ^RF^**  **([45.30 N, 132.57 E](https://goo.gl/maps/yFafyroN4RT2kYyPA))** | Xingkai Lake (Lake Khanka) is a large transboundary freshwater lake on the border between the PRC and the Russian Federation. The extensive floodplain wetlands around the lake provide stopover and breeding habitats for up to two million waterbirds, notably large congregations of Red-crowned Crane (VU) and Oriental Stork (EN). At least 14 waterbird species have internationally important populations in the national nature reserve ([RIS](https://rsis.ramsar.org/RISapp/files/RISrep/CN1155RIS_1708_en.pdf)) |
| **Sanjiang National Nature Reserve ^RF^**  **(**[**47.90 N, 134.50 E**](https://goo.gl/maps/cX57Y84RBivcmFx17)**)** | The Sanjiang Plain is an extensive delta alluvial plain where the Heilongjiang and Wusuli (Ussuri) Rivers converge. It supports 50,000-100,000 geese and ducks during the migration seasons, including up to 24,000 Swan Goose (VU), and also supports breeding populations of Red-crowned Crane (VU), White-naped Crane (VU) and Oriental Stork (EN) ([RIS](https://rsis.ramsar.org/RISapp/files/RISrep/CN1152RIS.pdf)). |

**Jilin Province**

| **SITE** | **IMPORTANCE OF SITE FOR MIGRATORY WATERBIRDS** |
| --- | --- |
| **Momoge National Nature Reserve ^R^**  **([46.00 N, 123.75 E](https://goo.gl/maps/yR2ioNPsBSvfKYQG8))** | Momoge NNR comprises extensive marshlands, grasslands and agricultural land that support more than 100,000 waterbirds during the migration seasons, including Anatidae, cranes and Oriental Stork (EN). Most notably, approximately 95% of the global population of Siberian Crane (CR) stops over there on migration (Wang *et al.* 2022). |
| **Xianghai National Nature Reserve ^RF^**  **([45.08 N, 122.33 E](https://goo.gl/maps/K44YaUgDv5tTwoHD8))** | Xianghai NNR is located at the western edge of the Songnen Plain, and the numerous lakes and extensive marshlands there are an important breeding and staging area for migratory waterbirds, including Anatidae, cranes and Oriental Stork (EN) ([RIS](https://rsis.ramsar.org/RISapp/files/RISrep/CN548RIS.pdf)). |

**Inner Mongolia**

| **SITE** | **IMPORTANCE OF SITE FOR MIGRATORY WATERBIRDS** |
| --- | --- |
| **Dalai Lake National Nature Reserve ^RF^**  **([49.00 N, 117.33 E](https://goo.gl/maps/HwfK1zJXc1wxubos6))** | Dalai Hu is one of the five largest freshwater lakes (by area) in the PRC and is surrounded by vast areas of marshland and arid steppes. Hundreds of thousands of waterbirds stage or breed there every year, including at least 11 species that occur in internationally important numbers ([RIS](https://rsis.ramsar.org/RISapp/files/RISrep/CN1146RIS.pdf), [SIS](https://eaaflyway.net/wp-content/uploads/2017/12/SIS-EAAF064-Dalai-Hu-National-Nature-Reserve_v2017.pdf)). |
| **Wuliangsuhai National Nature Reserve**  **(**[**41.10 N, 108.83 E**](https://goo.gl/maps/ynPoLMupp1jDGYC3A)**)** | Wuliangsuhai is a large lake with extensive emergent vegetation, that is an important breeding and staging area for waterbirds in a vast arid region of northwest PRC. A waterbird survey in 2011-2012 counted more than 165,900 individual birds there, including 19 species that exceeded 1% of flyway population thresholds, notably Swan Goose (VU), Far Eastern Curlew (EN) and Relict Gull (VU) (Zhang *et al.* 2017). |

***Yellow River Basin***

**Hebei**

| **SITE** | **IMPORTANCE OF SITE FOR MIGRATORY WATERBIRDS** |
| --- | --- |
| **Hengshui Lake National Nature Reserve ^F^**  **([37.60 N, 115.60 E](https://goo.gl/maps/1h4bMEn1o8awxodo9))** | Hengshui Lake NNR is an important breeding and stopover site for waterbirds in the arid North China Plain region. Nineteen waterbird species have occurred there in internationally important numbers, most notably around 15% of the global population of Baer’s Pochard (CR) and 30% of the flyway population of Bean Goose (Guo *et al.* 2021). |

**Qinghai Province**

| **SITE** | **IMPORTANCE OF SITE FOR MIGRATORY WATERBIRDS** |
| --- | --- |
| **Zhaling Lake-Er’ling Lake National Nature Reserve ^R^**  **(**[**34.93 N, 97.45 E**](https://goo.gl/maps/Uu7ZCaHkWk847fBX7)**)** | Zhaling Lake-Er’ling Lake NNR includes two high-altitude lakes that are important for migratory waterbirds, including at least five species that exceed 1% of flyway population thresholds there, notably Black-necked Crane (VU) (Sun *et al.* 2020, Gong *et al.* 2022). |

**Gansu Province**

| **SITE** | **IMPORTANCE OF SITE FOR MIGRATORY WATERBIRDS** |
| --- | --- |
| **Gahai-Zecha National Nature Reserve ^R^**  **([34.23 N, 102.32 E](https://goo.gl/maps/zjdGmoyh9qTfAkQe6))** | Gahai-Zecha NNR is a high-altitude wetland in arid Gansu Province, where at least four migratory waterbird species exceed 1% of flyway population thresholds, notably a large breeding population of Common Redshank and Black-necked Crane (VU) ([RIS](https://rsis.ramsar.org/RISapp/files/RISrep/CN1975RIS.pdf)). |

**Ningxia Province**

| **SITE** | **IMPORTANCE OF SITE FOR MIGRATORY WATERBIRDS** |
| --- | --- |
| **Shahu Lake National Wetland Park**  **(**[**38.82 N, 106.35 E**](https://goo.gl/maps/K3BPJvYQABUCJyzt8)**)** | Shahu Lake National Wetland Park is one of the few extensive wetlands in arid Ningxia Province. Based on the habitats present it is considered likely to support significant populations of migratory waterbirds, but no surveys are known to have been conducted to investigate their status. |

**Shaanxi Province**

| **SITE** | **IMPORTANCE OF SITE FOR MIGRATORY WATERBIRDS** |
| --- | --- |
| **Hongjianlao National Nature Reserve**  **(**[**39.08 N, 109.92 E**](https://goo.gl/maps/SHQneHmQhYyeVFXD8)**)** | Hongjianlao NNR is a wetland in an arid region of Shaanxi Province that in recent years has supported more than half of the global population of Relict Gull (VU), which nests on islands in the lake. At least three other migratory waterbird species exceed 1% of flyway population thresholds there, but the wetlands are subject to shrinkage linked to water extraction (Liang & Yan 2017, Liu *et al.* 2017). |

**Shanxi Province**

| **SITE** | **IMPORTANCE OF SITE FOR MIGRATORY WATERBIRDS** |
| --- | --- |
| **Yuncheng Yellow River Wetland Provincial Nature Reserve**  **(**[**35.03 N, 110.98 E**](https://goo.gl/maps/vTRhom9DWpqwPTU3A)**)** | Yuncheng Yellow River Wetland Provincial NR is the largest wetland nature reserve in Shanxi Province. It supports an important population of Whooper Swan, with more than 10,000 individuals estimated to spend the non-breeding period, and several other migratory waterbird species have exceeded 1% of flyway population thresholds there. |

**Henan Province**

| **SITE** | **IMPORTANCE OF SITE FOR MIGRATORY WATERBIRDS** |
| --- | --- |
| **Yellow River Wetland National Nature Reserve**  **(**[**34.87 N, 112.50 E**](https://goo.gl/maps/s3T8TAiBJuT9tuv37)**)** | Yellow River Wetland NNR is an extensive area of riverine floodplain wetlands that is a demonstration site for ADB’s Yellow River Project. It supports a large non-breeding population of Whooper Swan, and several other migratory waterbird species have exceeded 1% of flyway population thresholds there. |
| **Minquan Yellow River Old Riverway National Wetland Park ^R^ (**[**34.65 N, 115.32E**](https://goo.gl/maps/92xXzuUDhregiBEN6)**)** | Minquan Yellow River Old Riverway National Wetland Park comprises wetlands in the old channel of the Yellow River and is important for the conservation of migratory waterbirds (Li Changkan *et al.* 2019). Waterbird surveys in 2017-2019 counted up to 60,000 individual birds there, including six species that exceeded 1% of flyway population thresholds, most notably around 12.5% of the global population of Baer’s Pochard (CR) ([RIS](https://rsis.ramsar.org/RISapp/files/RISrep/CN2426RIS_2008_en.pdf)). |

***Yangtze Basin***

**Jiangxi Province**

| **SITE** | **EAAF SPECIES (≥1%)** | **SCORE** |
| --- | --- | --- |
| **Poyang Lake National Nature Reserve ^RF^**  **(**[**29.17 N, 115.97 E**](https://goo.gl/maps/hSAZ16nN5Q32ECfL9)**)** | Tundra Swan *Cygnus columbianus* *  Swan Goose *Anser cygnoid* (VU)**  Bean Goose *Anser fabalis* **  Greater White-fronted Goose *Anser albifrons* **  Lesser White-fronted Goose *Anser erythropus* (VU)  Falcated Duck *Mareca falcata*  Great Crested Grebe *Podiceps cristatus*  Siberian Crane *Leucogeranus leucogeranus* (CR)**  White-naped Crane *Grus vipio* (VU)*  Red-crowned Crane *Grus japonensis* (VU)**  Common Crane *Grus grus* *  Hooded Crane *Grus monacha* (VU)*  Black Stork *Ciconia nigra*  Oriental Stork *Ciconia boyciana* (EN)*  Eurasian Spoonbill *Platalea leucorodia* *  Pied Avocet *Recurvirostra avosetta* *  Black-tailed Godwit *Limosa limosa* **  Spotted Redshank *Tringa erythropus* | 825.49 |
| **Nanjishan Wetland Nature Reserve ^RF^**  **(**[**29.00 N, 116.28 E**](https://goo.gl/maps/z1FLTGbaW1NueGUeA)**)** | Tundra Swan *Cygnus columbianus*  Swan Goose *Anser cygnoid* (VU)  Bean Goose *Anser fabalis* *  Greater White-fronted Goose *Anser albifrons* *  Great Crested Grebe *Podiceps cristatus*  Siberian Crane *Leucogeranus leucogeranus* (CR)  White-naped Crane *Grus vipio* (VU)**  Common Crane *Grus grus*  Hooded Crane *Grus monacha* (VU)  Oriental Stork *Ciconia boyciana* (EN)  Eurasian Spoonbill *Platalea leucorodia*  Pied Avocet *Recurvirostra avosetta*  Black-tailed Godwit *Limosa limosa*  Spotted Redshank *Tringa erythropus* | 134.9 |

**Hunan Province**

| **SITE** | **EAAF SPECIES (≥1%)** | **SCORE** |
| --- | --- | --- |
| **East Dongting Lake National Nature Reserve ^R^**  **(**[**29.33 N, 112.92 E**](https://goo.gl/maps/wf5taALPX9xcgfcA7)**)** | Tundra Swan *Cygnus columbianus*  Greylag Goose *Anser anser*  Swan Goose *Anser cygnoid* (VU) *  Bean Goose *Anser fabalis* *  Greater White-fronted Goose *Anser albifrons* *  Lesser White-fronted Goose *Anser erythropus* (VU) *  Ruddy Shelduck *Tadorna ferruginea*  Falcated Duck *Mareca falcata**  Black Stork *Ciconia nigra*  Eurasian Spoonbill *Platalea leucorodia* *  Great White Egret *Ardea alba*  Dalmatian Pelican *Pelecanus crispus*  Great Cormorant *Phalacrocorax carbo*  Pied Avocet *Recurvirostra avosetta*  Spotted Redshank *Tringa erythropus* | 134.53 |
| **West Dongting Lake National Nature Reserve ^R^**  **(**[**28.82 N, 112.20 E**](https://goo.gl/maps/Msf2BW5KGB7hWGnG8)**)** | Tundra Swan *Cygnus columbianus*  Falcated Duck *Mareca falcata*  Eurasian Spoonbill *Platalea leucorodia*  Great Cormorant *Phalacrocorax carbo* | 13.88 |

**Anhui Province**

| **SITE** | **EAAF SPECIES (≥1%)** | **SCORE** |
| --- | --- | --- |
| **Shengjin Lake Nature Reserve ^RF^**  **(**[**30.38 N, 117.07 E**](https://goo.gl/maps/zA9LdVYQXj54GvEZ8)**)** | Swan Goose *Anser cygnoid* (VU)*  Bean Goose *Anser fabalis* *  Greater White-fronted Goose *Anser albifrons* *  Lesser White-fronted Goose *Anser erythropus* (VU)*  Falcated Duck *Mareca falcata*  Hooded Crane *Grus monacha* (VU)*  Eurasian Spoonbill *Platalea leucorodia*  Dalmatian Pelican *Pelecanus crispus*  Great Cormorant *Phalacrocorax carbo* | 99.59 |
| **Anqing Yangtze Riverine Wetland Nature Reserve ^F^**  **(**[**30.53 N, 116.90 E**](https://goo.gl/maps/kfAVnjWtK3LCuNQQA)**)** | Tundra Swan *Cygnus columbianus* *  Swan Goose *Anser cygnoid* (VU)  Bean Goose *Anser fabalis*  Greater White-fronted Goose *Anser albifrons*  Lesser White-fronted Goose *Anser erythropus* (VU)  Smew *Mergellus albellus*  Great Crested Grebe *Podiceps cristatus*  Hooded Crane *Grus monacha* (VU)  Oriental Stork *Ciconia boyciana* (EN)  Eurasian Spoonbill *Platalea leucorodia*  Great Cormorant *Phalacrocorax carbo* | 71.32 |

**Hubei Province**

| **SITE** | **EAAF SPECIES (≥1%)** | **SCORE** |
| --- | --- | --- |
| **Wanghu Lake National Nature Reserve ^R^**  **([29.87 N, 115.33 E](https://goo.gl/maps/mWu8pdanijCT9pWG6))** | Tundra Swan *Cygnus columbianus*  Bean Goose *Anser fabalis*  Great Crested Grebe *Podiceps cristatus*  Black Stork *Ciconia nigra*  Eurasian Spoonbill *Platalea leucorodia*  Great Cormorant *Phalacrocorax carbo* | 21.11 |
| **Chenhu Lake Provincial Nature Reserve ^R^**  **([30.30 N, 113.82 E](https://goo.gl/maps/HV1goGFc7TEHfa2LA))** | Bean Goose *Anser fabalis*  Falcated Duck *Mareca falcata*  Eurasian Spoonbill *Platalea leucorodia*  Dalmatian Pelican *Pelecanus crispus*  Great Cormorant *Phalacrocorax carbo* | 13.92 |

* Exceeding 10% of CSR1 estimate

** Exceeding 50% of CSR1 estimate

Sites in **bold** overlap with the protected area(s)

***Indonesia***

| **SITE** | **EAAF SPECIES (≥1%)** | **SCORE** |
| --- | --- | --- |
| Batu Bara coast, North Sumatra  **(**[**3.38 N, 99.43 E**](https://goo.gl/maps/ajna8QL4eECz13Gv9)**)** | Asian Dowitcher *Limnodromus semipalmatus*  Lesser Sandplover *Charadrius mongolus*  Spotted Greenshank *Tringa guttifer* (EN)  Black-tailed Godwit *Limosa limosa*  Terek Sandpiper *Xenus cinereus*  Common Redshank *Tringa totanus*  Curlew Sandpiper *Calidris ferruginea*  Broad-billed Sandpiper *Calidris falcinellus*  Bar-tailed Godwit *Limosa lapponica*  Eurasian Curlew *Numenius arquata*  Whimbrel *Numenius phaeopus*  Pacific Golden Plover *Pluvialis fulva*  Ruddy Turnstone *Arenaria interpres* ^1^ | 44.9 |
| **Banyuasin Peninsula and Berbak-Sembilang National Park** ^RF^, South Sumatra  **(**[**-1.89 N, 104.61 E**](https://goo.gl/maps/5KB9YTQngGp6qeATA)**)** | Black-tailed Godwit *Limosa limosa*  Lesser Sandplover *Charadrius mongolus*  Bar-tailed Godwit *Limosa lapponica*  Asian Dowitcher *Limnodromus semipalmatus*  Far Eastern Curlew *Numenius madagascariensis* (EN) ^1^ | 27.1 |
| **Deli-Serdang**, North Sumatra  **(**[**3.73 N, 98.79 E**](https://goo.gl/maps/utMbfK9ameTrZBm59)**)** | Lesser Sandplover *Charadrius mongolus*  Eurasian Curlew *Numenius arquata*  Spotted Greenshank *Tringa guttifer* (EN)  Ruff *Calidris pugnax*  Red Knot *Calidris canutus*  Ruddy Turnstone *Arenaria interpres*  Whimbrel *Numenius phaeopus* | 17.4 |
| **Wasur National Park** ^RF^, South Papua  **(**[**-8.60 N, 140.80 E**](https://goo.gl/maps/eDCmQe6KPcwU9X3R9)**)** | Lesser Sandplover *Charadrius mongolus* * | 10.4 |
| **Middle Mahakam Lakes**, East Kalimantan  **(**[**-0.36 N, 116.20 E**](https://goo.gl/maps/vENXFwx2vFciHKsz6)**)** | Wood Sandpiper *Tringa glareola* | 7.7 |
| **Sungai Progo Delta - Trisik Beach**, Yogyakarta  ([-7.98 N, 110.20 E](https://goo.gl/maps/F3v9AL1pwgKn9JUD8)) | Sanderling *Calidris alba*  Wood Sandpiper *Tringa glareola* | 7.4 |
| **Cemara Beach**, Jambi  **(**[**-1.42 N, 104.46 E**](https://goo.gl/maps/Z9CKK5ZGjpqYG4UCA)**)** | Spotted Greenshank *Tringa guttifer* (EN)  Lesser Sandplover *Charadrius mongolus*  Black-tailed Godwit *Limosa limosa*  Bar-tailed Godwit *Limosa lapponica*  Asian Dowitcher *Limnodromus semipalmatus* ^1^ | 7.1 |
| **East Aceh coast, Aceh**  **(**[**4.68 N, 97.96 E**](https://goo.gl/maps/DbjAnbv1m62fZMHo7)**)** | Lesser Sandplover *Charadrius mongolus* | 5.3 |
| **North Aceh coast, Aceh**  **(**[**5.22 N, 97.45 E**](https://goo.gl/maps/RT2WtrZ51hWPLDTb9)**)** | Lesser Sandplover *Charadrius mongolus*  Broad-billed Sandpiper *Calidris falcinellus* | 4.7 |
| **Kupang Bay, East Nusa Tenggara**  ([-10.05 N, 123.78 E](https://goo.gl/maps/1PT4dr1dqrN2JHG9A)) | Australian Pratincole *Stiltia isabella* | 3.5 |
| **Sekopong Bay, Way Kambas National Park**, Lampung  **(**[**-4.90 N, 105.88 E**](https://goo.gl/maps/Fk4Q4LxRKBmSUeer8)**)** | Asian Dowitcher *Limnodromus semipalmatus* | 2.2 |
| **Benoa Bay – Serangan Island**, Bali  **(**[**-8.74 N, 115.22 E**](https://goo.gl/maps/e5FUtoKKmRSamxuP6)**)** | Whimbrel *Numenius phaeopus* | 2.1 |
| **Gresik and Surabaya coast (incl. Ujung Pangkah)**, East Java  **(**[**-6.92 N, 112.62 E**](https://goo.gl/maps/3SgFXspYUVaM5MWi7)**)** | Red Knot *Calidris canutus* | 1.9 |
| Asahan coast, North Sumatra  **(**[**3.03N, 99.87 E**](https://goo.gl/maps/Q2tD9MPvAjdGA4Vj6)**)** | Lesser Sandplover *Charadrius mongolus* | 1.4 |
| North Labuhan Batu coast, North Sumatra  **(**[**2.77 N, 99.99 E**](https://goo.gl/maps/vZwgdJppStfpWa8u6)**)** | Common Redshank *Tringa totanus* | 1.4 |
| Labuhan Batu coast, North Sumatra  **(**[**2.70 N, 100.16 E**](https://goo.gl/maps/oaWUCVc2T52q54b78)**)** | Lesser Sandplover *Charadrius mongolus* | 1.4 |
| **North Seram coast (partly in Manusela National Park)**, Maluku  ([-2.95 N, 129.12 E](https://goo.gl/maps/uaTuxHsm9hZGpkoi9)) | Chinese Crested Tern *Thalasseus bernsteini* (CR) | 1.0 |

^1^ Nearly meeting 1% of CSR1 estimate

* Exceeding 10% of CSR1 estimate

Sites in **bold** overlap with the protected area(s)

***Lao People’s Democratic Republic***

| **SITE** | **EAAF SPECIES (≥1%)** | **SCORE** |
| --- | --- | --- |
| **Xe Pian NBCA (including Bueng Kiat Ngong wetlands)**, Champasak  **(**[**14.56 N, 106.06 E**](https://goo.gl/maps/y6j5k3LUh4XXXvCy7)**)** | Masked Finfoot *Heliopais personatus* (CR)  Asian Openbill *Anastomus oscitans* | 4.5 |
| Mekong River from Luang Prabang to Vientiane, Vientiane-Xayabury  **(**[**18.32 N, 101.52 E**](https://goo.gl/maps/4sb2nyuzANoLRA4y5)**)** | Little Ringed Plover *Charadrius dubius* | 2.2 |
| **Dong Kanthung Protected Forest (proposed NBCA)**, Champasak  **(**[**14.40 N, 105.43 E**](https://goo.gl/maps/WJStZZ1tM5hczr5B6)**)** | Masked Finfoot *Heliopais personatus* (CR) | 2.0 |

Sites in **bold** overlap with the protected area(s)

***Malaysia***

| **SITE** | **EAAF SPECIES (≥1%)** | **SCORE** |
| --- | --- | --- |
| **North-Central Selangor Coast**, Selangor  **(**[**3.24 N, 101.30 E**](https://goo.gl/maps/n9JhZ7VCU3zEGETM8)**)** | Lesser Sandplover *Charadrius mongolus*  Eurasian Curlew *Numenius arquata*  Common Redshank *Tringa totanus*  Whimbrel *Numenius phaeopus*  Spotted Greenshank *Tringa guttifer* (EN)  Painted Stork *Mycteria leucocephala* | 14.1 |
| Sadong-Saribas Coast, Sarawak  **(**[**1.68 N, 111.05 E**](https://goo.gl/maps/YQ93fXLMmBbXfrNe7)**)** | Lesser Sandplover *Charadrius mongolus*  Chinese Egret *Egretta eulophotes* (VU)  Terek Sandpiper *Xenus cinereus*  Whimbrel *Numenius phaeopus* | 13.2 |
| **Buntal Bay** ^F^, Sarawak  **(**[**1.69 N, 110.39 E**](https://goo.gl/maps/pAPhT8sH5JXxg2Ta7)**)** | Far Eastern Curlew *Numenius madagascariensis* (EN)  Lesser Sandplover *Charadrius mongolus*  Whimbrel *Numenius phaeopus*  Spotted Greenshank *Tringa guttifer* (EN)  Red Knot *Calidris canutus*  Gull-billed Tern *Gelochelidon nilotica* ^1^ | 9.2 |
| Teluk Air Tawar-Kuala Muda IBA, Pulau Pinang  **(**[**5.53 N, 100.37 E**](https://goo.gl/maps/s35e1Ltdj88JQQS5A)**)** | Lesser Sandplover *Charadrius mongolus*  Spotted Greenshank *Tringa guttifer* (EN)  Broad-billed Sandpiper *Calidris falcinellus* | 8.4 |
| Mersing-Endau Coast, Johor  **(**[**2.64 N, 103.71 E**](https://goo.gl/maps/ug4aivpN4ZbZ49dcA)**)** | Lesser Sandplover *Charadrius mongolus* | 2.2 |
| Pulau Bruit, Sarawak  **(**[**2.58 N, 111.29 E**](https://goo.gl/maps/gLTSeKf2hJq6pabc9)**)** | Chinese Egret *Egretta eulophotes* (VU) | 1.8 |

^1^ Nearly meeting 1% of CSR1 estimate

Sites in **bold** overlap with the protected area(s)

***Mongolia***

| **SITE** | **EAAF SPECIES (≥1%)** | **RANK** |
| --- | --- | --- |
| **Khar-Us Lake**  **(including Khovd River Tributary and Khar Lake)** ^RF^, Uvs Aimag  **(**[**47.75 N, 92.17 E**](https://goo.gl/maps/dpM1cC9thFRtc4pF6)**)** | Dalmatian Pelican *Pelecanus crispus* **  Great Crested Grebe *Podiceps cristatus*  Great Cormorant *Phalacrocorax carbo*  Greylag Goose *Anser anser*  Bar-headed Goose *Anser indicus*  Ruddy Shelduck *Tadorna ferruginea*  Common Goldeneye *Bucephala clangul*  Common Merganser *Mergus merganser*  Common Crane *Grus grus*  Northern Lapwing *Vanellus vanellus*  Caspian Tern *Hydroprogne caspia*  Great Egret *Egretta alba*  Whooper Swan *Cygnus cygnus*  Common Shelduck *Tadorna tadorna*  Mallard *Anas platyrhynchos*  Green-winged Teal *Anas crecca*  Gadwall *Anas strepera*  Eurasian Wigeon *Anas penelope*  Northern Pintail *Anas acuta*  Northern Shoveler *Anas clypeata*  Red-crested Pochard *Netta rufina*  Common Pochard *Aythya ferina* (VU)  Tufted Duck *Aythya fuligula*  Temminck’s Stint *Calidris temminckii*  Great Black-headed Gull *Larus ichthyaetus*  White Spoonbill *Platalea leucorodia*  Swan Goose *Anser cygnoid* (VU) | 1 |
| **Khurkh Khuiten** ^RF^, Khentii Aimag  **(**[**48.32 N, 110.37 E**](https://goo.gl/maps/cJRc8QYZGm59DPbe7)**)** | Ruddy Shelduck *Tadorna ferruginea* **  White-naped Crane *Grus vipio* (VU)  Great Crested Grebe *Podiceps cristatus*  Black Stork *Ciconia nigra*  Bean Goose *Anser fabalis*  Swan Goose *Anser cygnoid* (VU)  Whooper Swan *Cygnus cygnus*  Common Crane *Grus grus*  Demoiselle Crane *Anthropoides virgo*  Northern Lapwing *Vanellus vanellus* | 2 |
| **Mongol Daguur IBA including Ulz River and Khukh Lake (include SPA)** ^RF^, Dornod Aimag  **(**[**49.72 N, 115.25 E**](https://goo.gl/maps/mVrEasNrVML8RPFf8)**)** | Swan Goose *Anser cygnoid* (VU)  Ruddy Shelduck *Tadorna ferruginea*  Hooded Crane *Grus monacha* *  Common Pochard *Aythya ferina* (VU)  Whooper Swan *Cygnus cygnus*  Swan Goose *Anser cygnoid* (VU)  Common Shelduck *Tadorna tadorna*  Pacific Golden Plover *Pluvialis fulva*  Spotted Redshank *Tringa erythropus*  Pied Avocet *Recurvirostra avosetta*  Great Crested Grebe *Podiceps cristatus*  Great Cormorant *Phalacrocorax carbo*  Bean Goose *Anser fabalis*  Common Crane *Grus grus*  White-naped Crane *Grus vipio* (VU)  Demoiselle Crane *Anthropoides virgo* | 3 |
| **Uvs Lake** ^RF^, Uvs Aimag  **(**[**50.2 N, 92.28 E**](https://goo.gl/maps/ifViyK9kU4rVfMkz5)**)** | Eurasian Spoonbill *Platalea leucorodia*  Great Crested Grebe *Podiceps cristatus*  Dalmatian Pelican *Pelecanus crispus*  Great Cormorant *Phalacrocorax carbo*  Greylag Goose *Anser anser*  Ruddy Shelduck *Tadorna ferruginea* | 4 |
| **Tolbo Lake**, Bayan-Olgii Aimag  **(**[**48.53 N, 90.1 E**](https://goo.gl/maps/mG6TH75fZGvtgco58)**)** | Great Cormorant *Phalacrocorax carbo*  Bar-headed Goose *Anser indicus*  Whooper Swan *Cygnus cygnus* | 5 |
| **Airag Lake** ^RF^, Uvs Aimag  **([48.9 N, 93.43 E](https://goo.gl/maps/dj6JK8MKriPQcqRB8))** | Great Cormorant *Phalacrocorax carbo*  Pallas's Gull *Larus ichthyaetus*  Great Crested Grebe *Podiceps cristatus*  Dalmatian Pelican *Pelecanus crispus*  Ruddy Shelduck *Tadorna ferruginea*  Common Goldeneye *Bucephala clangul*  Common Merganser *Mergus merganser*  Northern Lapwing *Vanellus vanellus* | 6 |
| Tashgain Tavan Lakes, Dornod Aimag  **(**[**47.37 N, 118.45 E**](https://goo.gl/maps/FWTvPK3uCkQfVHPi9)**)** | Great Crested Grebe *Podiceps cristatus*  Swan Goose *Anser cygnoid* (VU)  Ruddy Shelduck *Tadorna ferruginea*  Common Shelduck *Tadorna tadorna*  Gadwall *Anas strepera*  Common Pochard *Aythya ferina* (VU)  White-naped Crane *Grus vipio* (VU)  Demoiselle Crane *Anthropoides virgo* | 7 |
| **Terkhiin Tsagaan Lake** ^RF^, Arkhangai Aimag  **(**[**48.17 N, 99.75 E**](https://goo.gl/maps/SR4F3LzAc1Pk7rbK7)**)** | Great Cormorant *Phalacrocorax carbo*  Bar-headed Goose *Anser indicus*  Ruddy Shelduck *Tadorna ferruginea*  Common Goldeneye *Bucephala clangul*  Common Merganser *Mergus merganser*  Northern Lapwing *Vanellus vanellus* | 8 |
| **Ogii Lake** ^RF^, Arkhangai Aimag  **(**[**47.77 N, 102.7 E**](https://goo.gl/maps/KzeHyttb8JjUA6sn8)**)** | Swan Goose *Anser cygnoid* (VU)  Great Crested Grebe *Podiceps cristatus*  Bar-headed Goose *Anser indicus*  Whooper Swan *Cygnus cygnus*  Ruddy Shelduck *Tadorna ferruginea*  Common Goldeneye *Bucephala clangul*  Common Crane *Grus grus*  Northern Lapwing *Vanellus vanellus*  Mute Swan *Cygnus olor* | 9 |
| **Buir Lake** ^RF^, Dornod Aimag (transboundary to Hulun Lake, PRC)  **(**[**47.77 N, 117.8 E**](https://goo.gl/maps/ZgbuftRscvSFVdzQ7)**)** | Great Crested Grebe *Podiceps cristatus*  Great Cormorant *Phalacrocorax carbo*  Grey Heron *Ardea cinerea*  Swan Goose *Anser cygnoid* (VU) | 10 |
| **Valley of lakes (including Boon Tsagaan, Orog Lake and Taatsiin Tsagaan** ^R^, Bayan-Khongor Aimag  **(**[**45.3 N, 100.12 E**](https://goo.gl/maps/eYcYNfz7v2GH15ZSA)**)** | Mute Swan *Cygnus olor* *  Common Shelduck *Tadorna tadorna*  Ruddy Shelduck *Tadorna ferruginea*  Great Cormorant *Phalacrocorax carbo*  Great Crested Grebe *Podiceps cristatus*  Hooded Crane *Grus monachal* | 11 |

* Exceeding 10% of CSR1 estimate

** Exceeding 50% of CSR1 estimate

Sites in **bold** overlap with a protected area(s)

***Philippines***

| **SITE** | **EAAF SPECIES (≥1%)** | **SCORE** |
| --- | --- | --- |
| North Manila Bay (Bulacan) (includes Pampanga River East Bank-Santa Cruz-Caliligawan-Pamawaran-Navotas-Taliptip-Tanza)  **(**[**14.77 N, 120.79 E**](https://goo.gl/maps/kCiNR7EpUB3DCeeDA)**)** | Kentish Plover *Charadrius alexandrinus*  Pacific Golden Plover *Pluvialis fulva*  Lesser Sandplover *Charadrius mongolus*  Marsh Sandpiper *Tringa stagnatilis*  Whiskered Tern *Chlidonias hybrida*  Common Redshank *Tringa totanus*  Curlew Sandpiper *Calidris ferruginea*  Caspian Tern *Hydroprogne caspia* | 24.6 |
| **Olango Island Wildlife Sanctuary** ^RF^, Cebu  **(**[**10.25 N, 124.05 E**](https://goo.gl/maps/aGgjBg2ivSVHmqHZ6)**)** | Lesser Sandplover *Charadrius mongolus* *  Common Tern *Sterna hirundo*  Grey-tailed Tattler *Tringa brevipes* (NT)  Ruddy Turnstone *Arenaria interpres*  Kentish Plover *Charadrius alexandrinus*  Whimbrel *Numenius phaeopus*  Red-necked Stint *Calidris ruficollis* (NT) | 24.6 |
| **Bangrin Marine Protected Area**, Pangasinan  **(**[**16.25 N, 119.93 E**](https://goo.gl/maps/UzefCJGtHAjpz2VH6)**)** | Intermediate Egret *Ardea intermedia* *  Great White Egret *Ardea alba* | 21.1 |
| **Negros Occidental Coastal Wetlands Conservation Area (NOCWCA)** ^RF^, Negros Occidental  **(**[**10.27 N, 122.84 E**](https://goo.gl/maps/ysqWL6989PcyfaCH7)**)** | Little Egret *Egretta garzetta*  Black-tailed Godwit *Limosa limosa* (NT)  Kentish Plover *Charadrius alexandrinus*  Caspian Tern *Hydroprogne caspia*  Chinese Egret *Egretta eulophotes* (VU)  Broad-billed Sandpiper *Calidris falcinellus*  Great Knot *Calidris tenuirostris* (EN)  Pacific Golden Plover *Pluvialis fulva*  Intermediate Egret *Ardea intermedia* | 14.8 |
| Lake Mainit, Agusan del Norte  **(**[**9.46 N, 125.52 E**](https://goo.gl/maps/r28meL9UqMMFjbEi8)**)** | Tufted Duck *Aythya fuligula* * | 13.8 |
| **North Manila Bay (Pampanga)** ^R^ (includes Pampanga River Westbank-Sasmuan Pampanga Coastal Wetlands)  **(**[**14.83 N, 120.59 E**](https://goo.gl/maps/ZD86vvpYWg7S8CjV8)**)** | Kentish Plover *Charadrius alexandrinus*  Intermediate Egret *Ardea intermedia*  Whiskered Tern *Chlidonias hybrida*  Great White Egret *Ardea alba*  Black-headed Gull *Larus ridibundus* | 12.2 |
| **Agusan Marsh Wildlife Sanctuary** ^R^, Agusan del Sur  **(**[**8.31 N, 125.90 E**](https://goo.gl/maps/T8mjuRw5zs4wQd956)**)** | Intermediate Egret *Ardea intermedia* | 6.9 |
| Candaba Wetlands, Pampanga  **(**[**15.07 N, 120.88 E**](https://goo.gl/maps/i7aLUiuQ38z9rS556)**)** | Garganey *Spatula querquedula* | 5.2 |
| Kabasalan-Siay Wetland Area (also known as Sibugay Wetlands), Zamboanga-Sibugay  **(**[**7.71 N, 122.81 E**](https://goo.gl/maps/7Fjp1veJV4kfE4mn9)**)** | Great White Egret *Ardea alba*  Intermediate Egret *Ardea intermedia*  Far-eastern Curlew *Numenius madagascariensis** | 3.1 |
| Panabo Coast, Davao del Norte  **(**[**7.29 N, 125.70 E**](https://goo.gl/maps/DowvWEGGwVwZ7MBM9)**)** | Chinese Crested Tern *Thalasseus bernsteini* (CR) | 3.0 |
| **North Manila Bay (Bataan) – Balanga Wetlands Park- Pilar Wetlands**  **(**[**14.69 N, 120.57 E**](https://goo.gl/maps/LUBBT6iv6BpDPFsC7)**)** | Kentish Plover *Charadrius alexandrinus*  Great White Egret *Ardea alba* | 2.6 |
| **Tubbataha Reef Natural Park** ^RF^, Palawan  **(**[**8.89 N,** **120.06 E**](https://goo.gl/maps/mt6nW2dANWGbWxXe7)**)** | Greater Crested Tern *Thalasseus bergii*  Black Noddy *Anous minutus* | 1.5 |

* Exceeding 10% of CSR1 estimate

Sites in **bold** overlap with a protected area(s)

***Thailand***

| **SITE** | **EAAF SPECIES (≥1%)** | **SCORE** |
| --- | --- | --- |
| **Pak Thale-Laem Phak Bia coast** ^F^, Phetchaburi  **(**[**13.07 N, 100.07 E**](https://goo.gl/maps/Xo9mo23s62Eup2Yv7)**)** | Spotted Greenshank (EN) *Tringa guttifer* *  Lesser Sandplover *Charadrius mongolus*  Broad-billed Sandpiper *Calidris falcinellus*  Ruff *Calidris pugnax*  Black-tailed Godwit *Limosa limosa*  Great Knot *Calidris tenuirostris* (EN)  Caspian Tern *Hydroprogne caspia*  Curlew Sandpiper *Calidris ferruginea* ^1^ | 26.1 |
| **Bueng Boraphet**, Nakhon Sawan  **(**[**15.70 N, 100.24 E**](https://goo.gl/maps/Jwfg2ygsHEFt85uQ7)**)** | Glossy Ibis *Plegadis falcinellus* *  Asian Openbill *Anastomus oscitans*  Intermediate Egret *Ardea intermedia*  Great White Egret *Ardea alba*  Garganey *Spatula querquedula*  Black-winged Stilt *Himantopus himantopus* ^1^ | 24.6 |
| Pak Nam Prasae, Rayong  **(**[**12.70 N, 101.71 E**](https://goo.gl/maps/nGR6k5SbbCEEx51LA)**)** | Lesser Sandplover *Charadrius mongolus*  Spotted Greenshank *Tringa guttifer* (EN) | 6.1 |
| **Huai Chorakhe Mak Reservoir**, Buriram  **(**[**14.90 N, 103.03 E**](https://goo.gl/maps/mVRp4i2dKx7bU1Wz9)**)** | Sarus Crane *Grus antigone sharpii* (VU) | 5.0 |
| **Krabi River Mouth** ^RF^, Krabi  **(**[**8.02 N, 98.93 E**](https://goo.gl/maps/MEHeSUFFi61ZEXeX8)**)** | Lesser Sandplover *Charadrius mongolus*  Spotted Greenshank (EN) *Tringa guttifer* | 4.0 |
| Khlong Tamru (Bang Pakong), Chachoengsao  **(**[**13.48 N, 100.94 E**](https://goo.gl/maps/h9gWB4Yu4Jfx9zVX6)**)** | Lesser Sandplover *Charadrius mongolus*  Ruff *Calidris pugnax* | 3.5 |
| Khlong Yai, Trat  **(**[**11.78 N, 102.88 E**](https://goo.gl/maps/V38Wt4T1iGP4Q7qw8)**)** | Lesser Sandplover *Charadrius mongolus*  Spotted Greenshank *Tringa guttifer* (EN) | 2.8 |
| **Sanam Bin Reservoir**, Buriram  **(**[**14.64 N, 103.08 E**](https://goo.gl/maps/98vyPfWCif39d2gD6)**)** | Sarus Crane *Grus antigone sharpii* (VU) | 2.5 |
| **Ko Libong Non-Hunting Area**  **and Hat Chao Mai National Park** ^F^, Trang  **(**[**7.29 N, 99.44 E**](https://goo.gl/maps/3vUHmgLyDeLPZEQs6)**)** | Lesser Sandplover *Charadrius mongolus*  Spotted Greenshank *Tringa guttifer* (EN) | 2.2 |
| **Khok Kham salt pans** ^F^, Samut Sakhon  **(**[**13.51 N, 100.35 E**](https://goo.gl/maps/L8CLGZsM2UgVag1U8)**)** | Spotted Greenshank *Tringa guttifer* (EN) | 1.9 |
| **Phang-nga Bay** ^R^, Phang Nga  **(**[**8.32 N, 98.52 E**](https://goo.gl/maps/FwtFkMMyryK6nSg28)**)** | Lesser Sandplover *Charadrius mongolus* | 1.5 |
| Bang Pu coast and fish ponds, Samut Prakan  **(**[**13.50 N, 100.74 E**](https://goo.gl/maps/L1RMrKsshPNGWAKm9)**)** | Black-tailed Godwit *Limosa limosa* | 1.1 |

^1^ Nearly meeting 1% of CSR1 estimate

^*^ Exceeding 10% of CSR1 estimate

Sites in **bold** overlap with a protected area(s)

***Viet Nam***

| **SITE** | **EAAF SPECIES (≥1%)** | **SCORE** |
| --- | --- | --- |
| **Tram Chim National Park** ^RF^, Dong Thap  **(**[**10.72 N, 105.51 E**](https://goo.gl/maps/R8AKs9Cx3UB4QMjF8)**)** | Painted Stork *Mycteria leucocephala*  Asian Openbill *Anastomus oscitans*  Sarus Crane *Grus antigone sharpii* (VU) | 7.8 |
| **Thai Thuy (including Thai Binh Wetland Protected Area)**, Thai Binh  **(**[**20.61 N, 106.66 E**](https://goo.gl/maps/5YQwa7dYwdY3MihL8)**)** | Common Tern *Sterna hirundo*  Little Tern *Sterna albifrons* | 6.3 |
| **Can Gio (incl. Can Gio Biosphere Reserve, Ly Nhon & Can Thanh salt pans, and coastal flats)**, Ho Chi Minh City  ([10.39 N, 106.93 E](https://goo.gl/maps/BHiPffYYSe3fnKXQ6)) | Lesser Sandplover *Charadrius mongolus*  Painted Stork *Mycteria leucocephala* | 4.0 |
| **Xuan Thuy National Park** ^R^, Nam Dinh  **(**[**20.21 N, 106.55 E**](https://goo.gl/maps/NJedWvmEj5C7nPpa7)**)** | Spotted Greenshank *Tringa guttifer* (EN)  Black-faced Spoonbill *Platalea minor* (EN)^1^ | 2.8 |
| Binh Dai IBA, Ben Tre  **(**[**10.08 N, 106.76 E**](https://goo.gl/maps/Soyd2nX9iW2aYTqu5)**)** | Lesser Sandplover *Charadrius mongolus* | 2.7 |
| Tan Thanh-Con Ngang coast, Tien Giang  **(**[**10.26 N, 106.76 E**](https://goo.gl/maps/nnwHGzT6DTrJhUPq9)**)** | Lesser Sandplover *Charadrius mongolus* | 2.6 |
| Ba Tri IBA, Ben Tre  **(**[**10.01 N, 106.69 E**](https://goo.gl/maps/z1p8wdqRXmwcWvmr5)**)** | Lesser Sandplover *Charadrius mongolus* | 2.4 |
| Do Son coast, Hai Phong  **(**[**20.70 N, 106.77 E**](https://goo.gl/maps/Tu9evdTEye2GTUe69)**)** | Spotted Redshank *Tringa erythropus* (VU) | 1.8 |
| An Hai (Cat Hai) IBA, Hai Phong  **(**[**20.79 N, 106.78 E**](https://goo.gl/maps/SFwBUcQCsNrUrTyQ6)**)** | Broad-billed Sandpiper *Calidris falcinellus* | 1.3 |

^1^ Nearly meeting 1% of CSR1 estimate

Sites in **bold** overlap with a protected area(s)
